# De novo discovery of conserved gene clusters in microbial genomes with Spacedust

**DOI:** 10.1101/2024.10.02.616292

**Authors:** Ruoshi Zhang, Milot Mirdita, Johannes Söding

## Abstract

Metagenomics has revolutionized environmental and human-associated microbiome studies. However, the limited fraction of proteins with known biological process and molecular functions presents a major bottleneck. In prokaryotes and viruses, evolution favors keeping genes participating in the same biological processes co-localized as conserved gene clusters. Conversely, conservation of gene neighborhood indicates functional association. Spacedust is a tool for systematic, *de novo* discovery of conserved gene clusters. To find homologous protein matches it uses fast and sensitive structure comparison with Foldseek. Partially conserved clusters are detected using novel clustering and order conservation P-values. We demonstrate Spacedust’s sensitivity with an all-vs-all analysis of 1 308 bacterial genomes, identifying 72 843 conserved gene clusters containing 58% of the 4.2 million genes. It recovered 95% of antiviral defense system clusters annotated by a specialized tool. Spacedust’s high sensitivity and speed will facilitate the annotation of the huge numbers of sequenced bacterial, archaeal and viral genomes.

## Introduction

In the last decade, metagenomics has accelerated the pace of research into microbial ecology and human-associated microbiomes and into their intimate association with our health (1). Hundreds of thousands of microbial and viral genomes assembled from shotgun metagenomics permit the study of microbes and their interactions with each other and their environment (2–4). However, our ability to extract useful insights from such data is severely limited by the lack of functional information (5). Even in well-studied ecosystems such as the human gut, for around 40% of genes neither molecular function nor biological process are annotatable (2).

The standard approach for protein function annotation is by homology inference, that is, by sequence similarity search to find the best match in reference databases such as InterPro, KEGG orthologs, COGs, or SEED (6–8), and transferring the annotation if certain criteria are met (9–13). Earlier approaches relied on sequence-sequence search tools such as BLAST. However, function can remain conserved even at sequence identities much below 20%, which these approaches cannot detect (14). Therefore, modern approaches with increased sensitivity search with the query protein sequences through databases of profile hidden Markov models (HMMs) or sequence profiles. These are precomputed from multiple sequence alignments of protein family members with the same or similar functions. Many databases of orthologous families have been developed to automate the process of clustering orthologous protein sequences together (7, 15). This approach is motivated by the “orthology conjecture”, which states that orthologous sequences are more likely to be functionally related than paralogous ones, although the difference appears to actually be small (16, 17).

Integration of genomic context can improve the precision of ortholog clustering and functional annotation. Proteins do not work in isolation but cooperate with others in biological pathways. Evolution has a tendency to keep functionally associated genes closely together in prokaryotic and viral genomes. This can be a consequence of co-expressed genes sharing regulatory sequences or even forming part of the same transcription unit, an operon (18). Clustering also maximizes the chances of horizontal transfer of useful gene modules, and it minimizes disruptions of functionally associated genes by genomic recombination (19–21). Some methods exploit gene neighborhood conservation to increase specificity for identifying orthologs (22–25). Others used the “guilt by association” principle to find functionally associated genes (e.g. (26, 27)). Many methods have been designed for detecting a specific type of clusters, such as biosynthetic gene clusters (BGCs) (28–31), phage defense systems (32–34), virulence and antibiotic resistance factors (35, 36), or xenobiotic degradation pathways (37). Most search the protein sequences from the query genome against a pre-assembled database of profile HMMs representing protein families typically occurring in these clusters. They then apply heuristic rules for what constitutes a valid cluster match.

A few tools aim to find genomic neighborhoods similar to a query neighborhood (38–41). The sensitivity of these *de novo* cluster detection methods is limited severely by the use of sequence-sequence comparison tools such as BLAST or DIAMOND (42, 43), compounding their ability to detect all but closely related conserved clusters (44). They also do not scale up to more than a few hundred genomes in all-vs-all search mode, and some require strict conservation of gene order (colinearity). Some approaches find conserved clusters by first searching each of the genomes to be analyzed against a profile HMM database of orthologous groups, and then find clusters of genes with similar composition of orthologs (45, 46). While this type of approach has improved sensitivity over the first, it requires a reference database of orthologous groups and therefore excludes the many proteins from as yet unknown families (47).

Spacedust is a tool for systematic *de novo* discovery of conserved gene clusters across multiple genomes. It finds all gene clusters significantly conserved between any two genomes in a set of input genomes. Conserved clusters are found by maximizing the statistical significance measured with two novel statistics assessing the degree of clustering and the degree of order and strand conservation. Since remote homologies are critical to achieve high sensitivity for detecting conserved gene clusters, Spacedust performs its homology searches with our novel structure-based search tool Foldseek. Foldseek has similar sensitivity as the best structural comparison tools and much higher than sequence-sequence-, sequence-profile- and profile-HMM-based searches. Foldseek has been shown to be much more sensitive than sequence-sequence-, sequence-profile- and profile-HMM-based searches (48, 49).

Spacedust improves upon previous methods in several ways: (i) It is reference-free and can discover conserved clusters of any type and composition, (ii) its structure-based search maximizes its sensitivity for finding remotely related conserved gene clusters, (iii) its high speed allows for analyzing a large number of genomes for conserved gene clusters using all-versus-all searches, (iv) it integrates functional annotation of proteins to facilitate inference of function from cluster members, and (v) it offers a user-friendly Google Colab notebook.

We demonstrate the utility of Spacedust by detecting conserved clusters in an all-versus-all comparison of 1 308 representative bacterial genomes from different genera with a total of 4.2 million protein-coding genes. Spacedust recovers previously annotated gene clusters, e.g., operons, antiviral defense systems, and BGCs. It is able to assign 58 % of all 4.2 M genes and 35% of genes without any annotation to conserved gene clusters. Spacedust also discovers the vast majority of antiphage defense systems in this data set and achieves better results in identifying 207 manually annotated biosynthetic gene clusters than three specialized tools.

## Results

### Spacedust Algorithm

Spacedust takes as input a set *Q* of query genomes and a set *T* of target genomes (which may be equal to *Q*) and, for each pair (*q, t*) ∈ *Q* ×*T* of query and target genome, it finds all gene clusters whose gene arrangement is at least partially conserved between *q* and *t* (Fig. 1 and Methods). For that purpose, Spacedust first identifies homologous matches (“hits”) between proteins in *Q* and proteins in *T* using our sensitive structure search tool Foldseek (48) and our sequence search tool MMseqs2 (50) (steps 1-5 in Fig. 1A). It then finds cluster matches between proteins clustered near each other in both the query and the target genomes using a greedy cluster detection algorithm (step 6 in Fig. 1A and Fig. 1B).

**Fig. 1.**
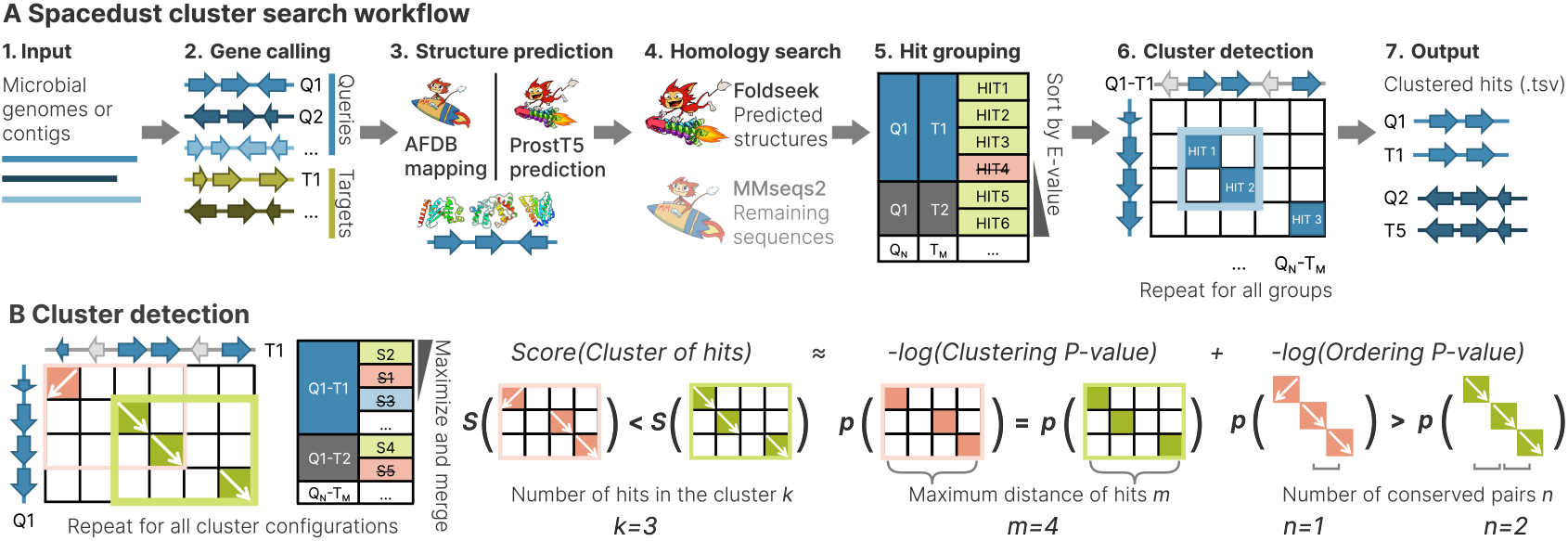
Spacedust algorithm. (A) **1** Input are microbial query and target genomes or assembled metagenomic contigs, (*Q*1,…), (*T* 1,…). **2** Predict protein-coding genes from all genomes *Q* and *T*. **3** Map protein sequences to AlphaFold DB (AFDB) using MMseqs2, or predict structure sequences with ProstT5. **4** Search all query proteins against all target proteins using Foldseek and MMseqs2, and combine search results. **5** Group hits between *Q − T* pairs, for each query sequence in *Q* select the hit in *T* with the best E-value. **6** For every *Q − T* pair, detect clusters of hits with significant conservation of gene neighborhood. **7** Report pairwise clusters of hits between *Q − T* in a tab-delimited file. (B) Starting from single-gene clusters, clusters are iteratively merged if this increases their score until all scores are maximal. Conservation of gene neighborhood for any given cluster of hits is evaluated by combining a clustering P-value and an ordering P-value.

Briefly, the algorithm starts with each protein hit in its own cluster and tries to add protein hits to the cluster matches one at a time. If the significance score of the cluster match improves, the addition is accepted and the algorithm continues, until the significance of the cluster matches cannot be improved further. The significance score is calculated as the sum of the negative logarithms of a clustering P-value and an ordering P-value. The clustering P-value is the probability of finding *by chance* at least *k* matches within a window of at most *m* genes in both the query and the target genome. The ordering P-value is the probability to find *by chance* at least *n* pairs of genes of the cluster match in conserved order in both genomes. The cluster detection algorithm thereby identifies positionally conserved clusters between all *Q* − *T* pairs of genomes. Optionally, the cluster matches for each query genome, aggregated across multiple reference genomes, can be visualized as a measure of conservation strength.

### A reference set of remotely conserved bacterial gene clusters

Despite the availability of tens of thousands of complete bacterial genomes, the gene cluster conservation landscape has yet to be surveyed systematically. To address this gap, we curated a dataset of 1 308 bacterial reference genomes, covering a broad phylogenetic range. These genomes were selected such that they belong to different bacterial genera (Methods), to focus on detecting remote homology and globally conserved clusters across higher taxonomic ranks. This choice means species- and genus-specific gene clusters cannot be found in this analysis. We subjected all predicted genes (4.19 million) from the 1 308 genomes to an all-vs-all Foldseek+MMseqs2 search using Spacedust. The all-vs-all homology search and hit filtering process took 72 hours to complete on two servers with two 64-core AMD EPYC ROME 7742 CPUs each, or 150 ms per genome-genome comparison (Supplementary material), and the subsequent cluster detection required 51 minutes. The run time of Spacedust scales quadratically with the total number of genomes and proteins in the genomes, due to the all-vs-all search and cluster detection. The search yielded 321.2 million cluster hits in 106.6 million cluster matches, with an average of 3 genes per cluster match (Fig. 2A).

**Fig. 2.**
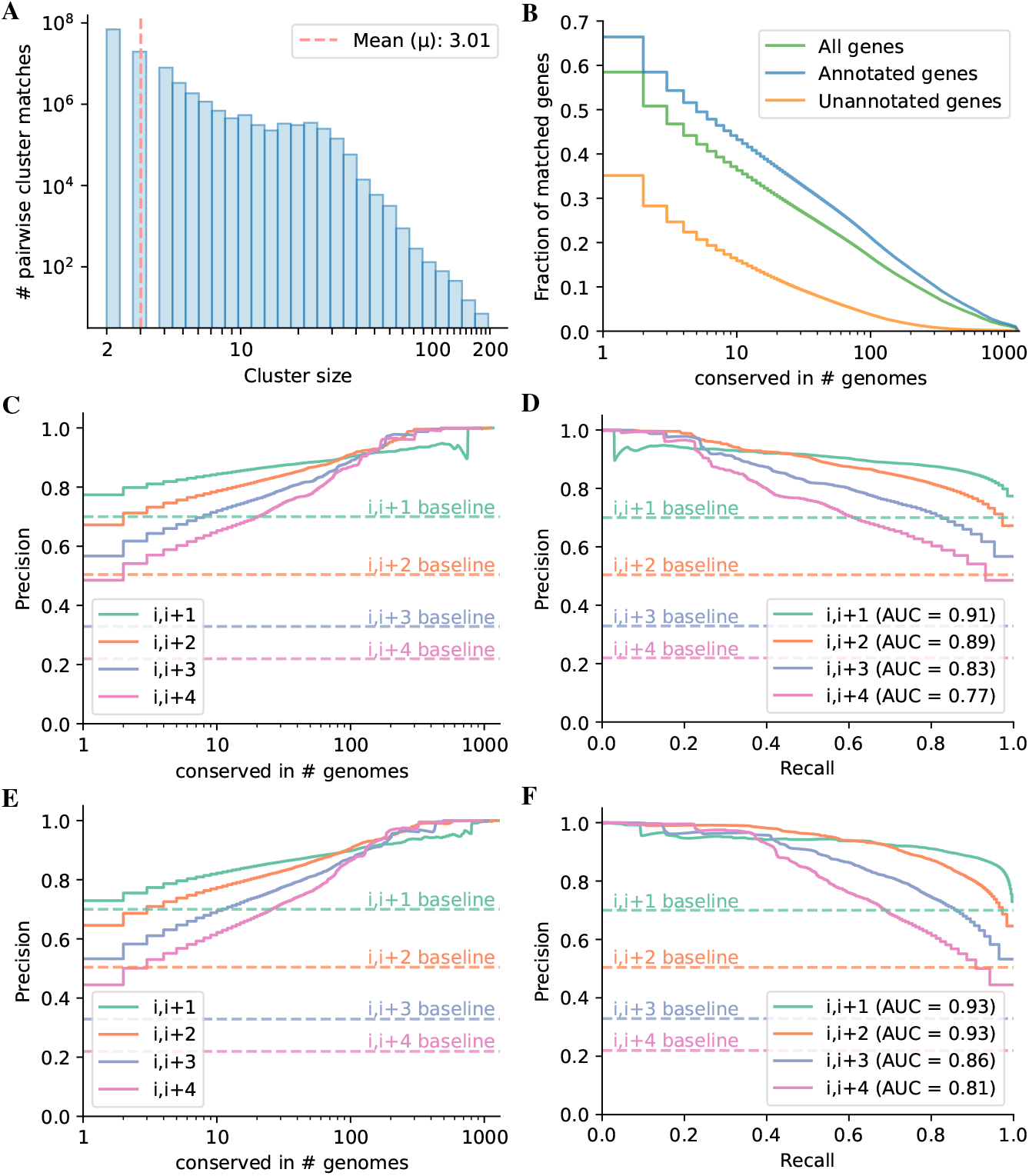
Conservation of gene clusters identified by Spacedust predicts functional association. (A) Distribution of cluster sizes of all 106.6 million pairwise cluster matches among 1 308 bacterial genomes. (B) Number of all (green), annotated (blue), and unannotated (orange) genes forming part of a cluster match in at least the number of genomes shown on the x-axis. (C, E) Precision of the functional association of gene pairs, separated by up to 4 genes in Spacedust cluster matches, versus the number of genomes in which the pair is conserved. True positive predictions are those gene pairs with the same KEGG module IDs. (C) uses Foldseek+MMseqs search, while (E) uses Foldseek-only search with ProstT5. (D, F) Precision versus recall of functional association of gene pairs separated by up to 4 genes. The analysis excludes ribosomal genes; see Fig. S7 with ribosomal genes. (D) uses Foldseek+MMseqs search, while (F) uses Foldseek-only search with ProstT5.

These pairwise cluster matches were subsequently grouped on the level of genome and genes, which yielded 72 483 non-redundant clusters comprising 2.45 million genes, representing 58% of the dataset. We classified 4.19 million genes based on their eggNOG-mapper annotations: “annotated” if the gene was assigned a specific function (3.13 million, or 75% of the dataset), or “unannotated” if the gene was labeled as “hypothetical protein”, “protein with unknown function”, or lacking any annotation (1.06 million, or 25% of the dataset). Notably, 66% of the annotated genes were found in non-redundant clusters present in more than one genome. Additionally, 35% of the unannotated genes were found in non-redundant clusters (Fig. 2B).

To evaluate the functional associations within the non-redundant clusters, we evaluated the congruence of KEGG module IDs for gapped gene pairs separated by up to four genes (*i, i* + 1) … (*i, i* + 4) (Fig. 2C, Fig. S7). A gene pair is considered a true positive if both genes share a common KEGG module ID, and false positive if not. The area under the precision-recall curve (PR-AUC) indicates that Spacedust identifies cluster matches with significantly higher accuracy than the baseline model, which assumes any neighboring gene pair (*i, i* + *x*) to be functionally associated. Similarly, we assessed the functional association within the non-redundant clusters predicted by the Foldseek-only search mode (Fig. 2E,F), with the PR-AUC for gapped gene pairs (*i, i* + 1) … (*i, i* + 4) slightly higher than that of the default Foldseek+MMseqs search mode.

### Global functional conservation of a Cyanobacterium genome

To illustrate how Spacedust can support functional annotation of genomes, we took one example genome of a unicellular cyanobacterium *Synechocystis* sp. PCC6803 from the reference database with the 1 308 genomes. This genome comprises one chromosome (GenBank accession: BA000022.2) and four plasmids (GenBank accession: AP004311.1, AP004312.1, AP004310.1, AP006585.1), totaling 3551 protein-coding genes. All the detected clusters are visualized as an interactive cluster heatmap (Fig. S1). For better visibility, we zoomed in on a specific region spanning protein location indices 500 to 800 (Fig 3A, corresponding to 0.0007% of the total dataset) and integrated functional annotation data obtained from eggNOG-mapper (51). This allowed us to assign functions to many of the proteins. From this selected genomic region containing 300 genes, we identified three distinct cyanobacteria-specific clusters, indicative of functional conservation across related species. Additionally, we detected 21 clusters shared with other phyla (Supplementary Table S2). Some clusters corresponded to single operons, while others spanned multiple operons.

**Fig. 3.**
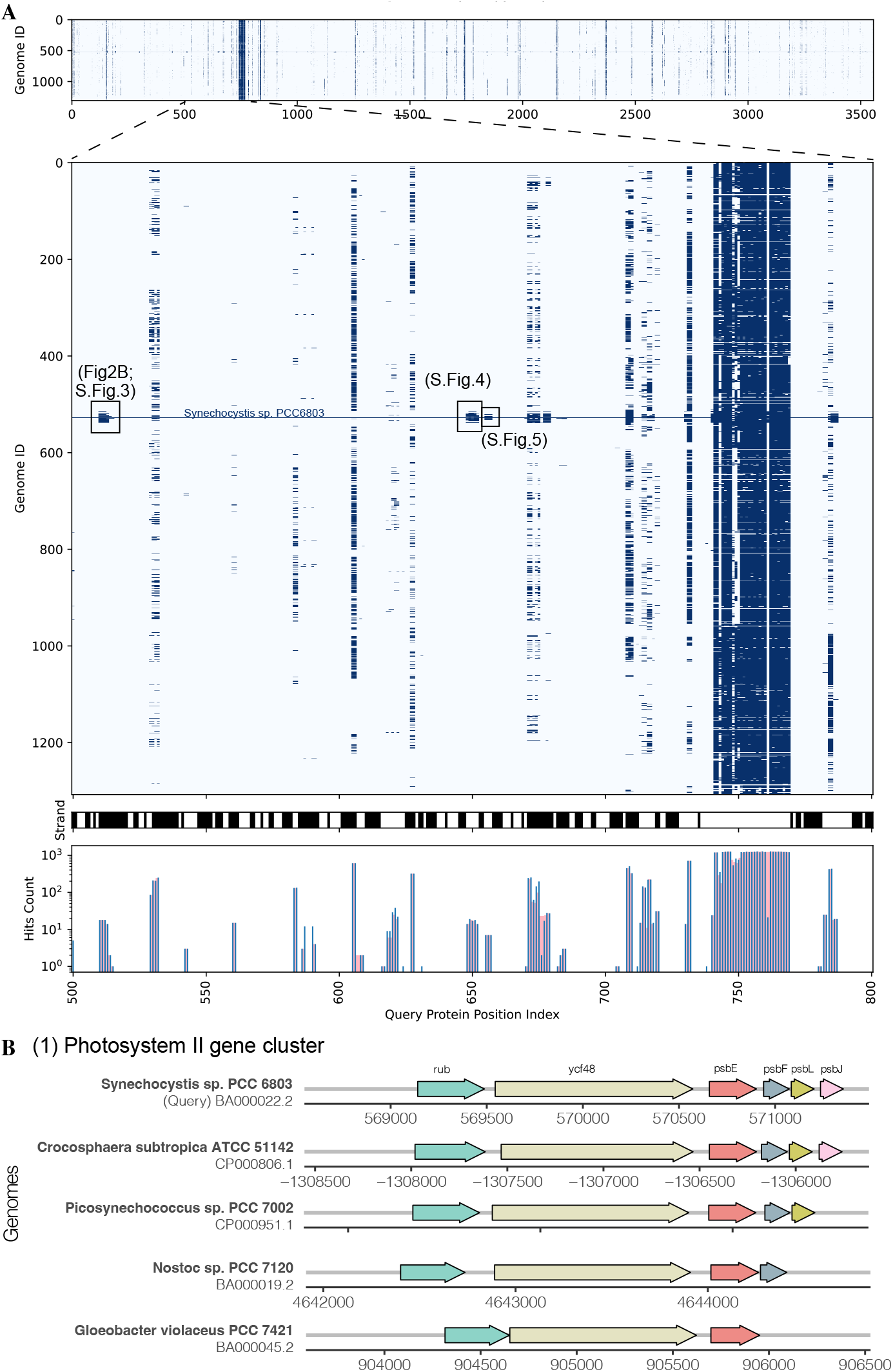
Evolutionary conservation of gene clusters in an example cyanobacterium. (A) Zoomed view of clustered hits of *Synechocystis* sp. PCC6803 (Genome ID 527) against 1 308 bacterial reference genomes. Query proteins with location indices 500-800 on the genome shown. Top panel: Cluster heatmap of the presence/absence of clustered hits across reference bacteria. Middle panel: Transcription direction (black: forward, white: reverse). Bottom panel: Number of clustered hits per protein (blue) and hit pairs in the same gene cluster (pink). (B) Example cyanobacteria-specific gene cluster 1. Gene names annotated by eggNOG-mapper are shown at the top.

Cluster 1 (Fig. 3B, Fig. S4) comprises genes associated with photosystem II (PSII), the protein-pigment complex that drives oxygenic photosynthesis. The first two genes, rubredoxin and *ycf48*, are crucial for PSII activity and assembly. The remaining genes, psbEFLJ, form an operon encoding components of the core PSII complex. In many cases, psbL and psbJ are absent, which could be owed to poor conservation or the short length of their sequences.

Cluster 2 (Fig. S5) forms an operon encompassing components of the phycobilisome complex rod, a large protein complex in cyanobacteria responsible for capturing sunlight and transferring energy to the photosynthetic reaction centers. The genes *cpcA* and *cpcB* encode two major subunits of the rod, while *cpcD, cpcC*, and *cpcC2* encode linker components connecting the rod to the PBS core (52). Conversely, homologous clusters in some other cyanobacteria only contain one copy of *cpcC* gene, suggesting that *cpcC* and *cpcC2* might have been created by gene duplication. In some genomes, the genes are still co-localized despite the order of the genes being only partially conserved.

In Cluster 3 (Fig. S6), the first two genes are both annotated as *spkA* by eggNOG-mapper, encoding a eukaryotic-type serine/threonine protein kinase involved in signal transduction and mobility. Alignment with other clusters revealed gene fusion of these two genes in other cyanobacteria (53). The third gene in the cluster is highly conserved in other genomes but could not be annotated using eggNOG-mapper.

### *De novo* identification of specialized gene clusters

To further assess Spacedust’s ability to identify conserved gene clusters, we focused on two categories of known, specialized gene clusters, antiviral defense systems and biosynthetic gene clusters (BGCs).

We used PADLOC v1.1.0 (33) to identify all known antiviral defense systems in the 1 308 bacterial reference genomes. We removed any predicted region consisting only of single genes. Spacedust was able to recover in total 5255 (95%) of 5520 multi-gene defense system clusters detected by PADLOC (Fig. 4A,B), with 93 % (4888) of the clusters matching fully and 7 % (367) matching partially to the PADLOC prediction. Most partial cluster matches resulted from missing matches to one or two short genes at the edge of longer clusters such as CRISPR-Cas systems. For 73 out of the 106 defense system types more than 90 % of all defense system clusters were discovered in their entirety by Spacedust (Fig. 4A), despite the fact that restriction-modification type II clusters are the most abundant type of defense systems in the dataset yet most challenging for Spacedust to detect.

**Fig. 4.**
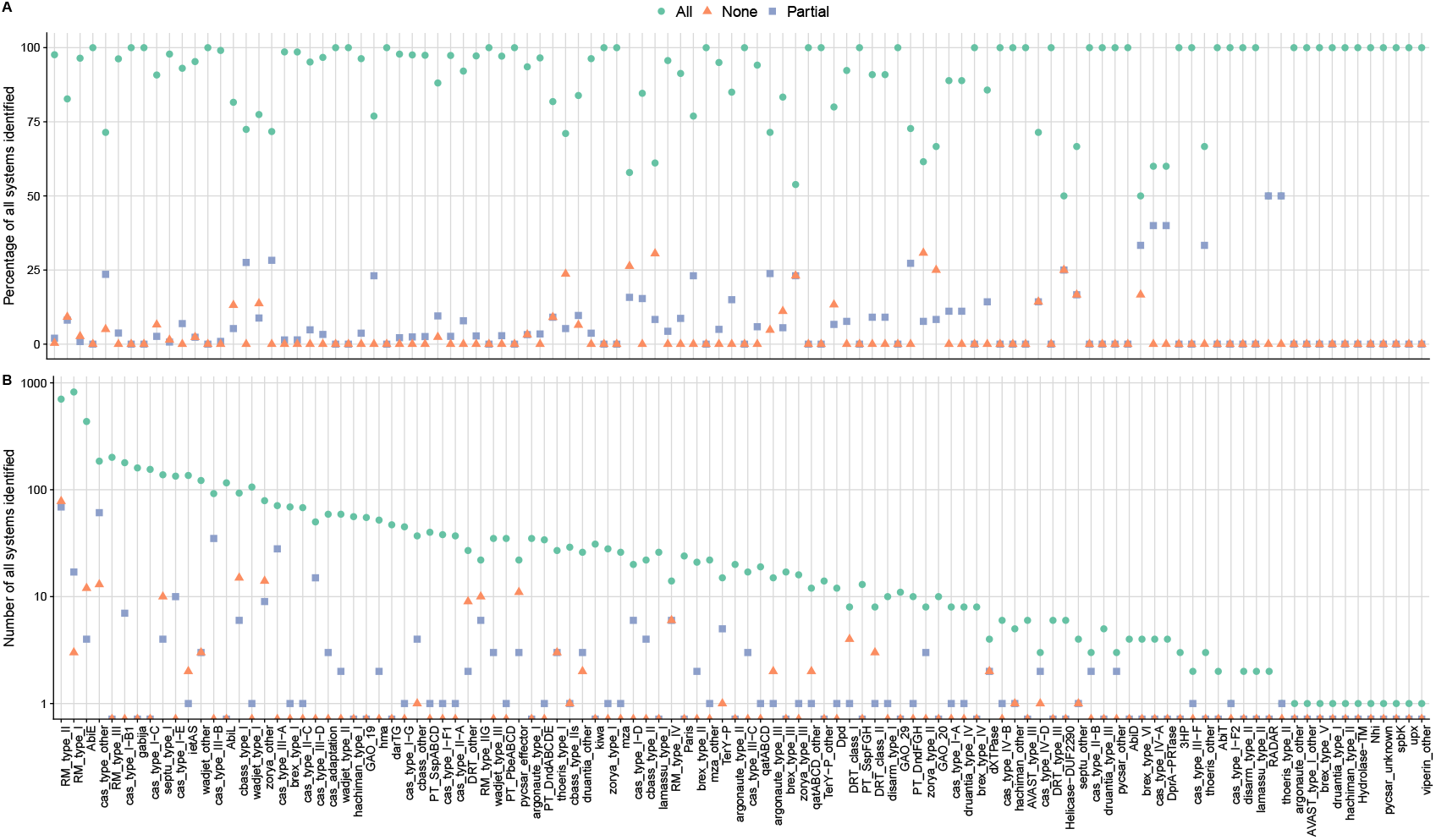
Spacedust recovers the vast majority of antiviral defense systems predicted by specialized tools. **(A)** Percentage and **(B)** number of multi-gene antiviral defense system clusters predicted by PADLOC which are also discovered by Spacedust in their entirety (green circle), partially (blue square), or missed (red triangle) within 1 308 bacterial genomes. 95 % of all defense system clusters predicted by PADLOC were recovered, 93 % of these in full length.

To evaluate Spacedust’s ability to recover biosynthetic gene clusters (BGCs), we utilized a gold standard dataset consisting of 9 complete genomes available at the NCBI that were fully annotated with BGC and non-BGC regions (29). We queried these 9 genomes against our reference set of 1 308 bacterial genomes using Spacedust. We compared the results with three tools specialized in identifying BGCs, ClusterFinder (29), DeepBGC (30) (using a cutoff of 10% false positive rate as recommended) and GECCO (31). As Spacedust returns all conserved clusters and not exclusively BGCs, it was not feasible to compare the precision of BGC detection based on all predictions. We therefore evaluated the F1 score, equal to the harmonic mean of precision and recall, for each of the annotated BGCs (Fig. 5). The precision is the fraction of genes in the overlapping predicted region that were annotated as BGC genes, and the recall is the fraction of genes in the annotated BGC region that were predicted as BGC region by the tool. Spacedust achieves higher F1 scores than ClusterFinder,

**Fig. 5.**
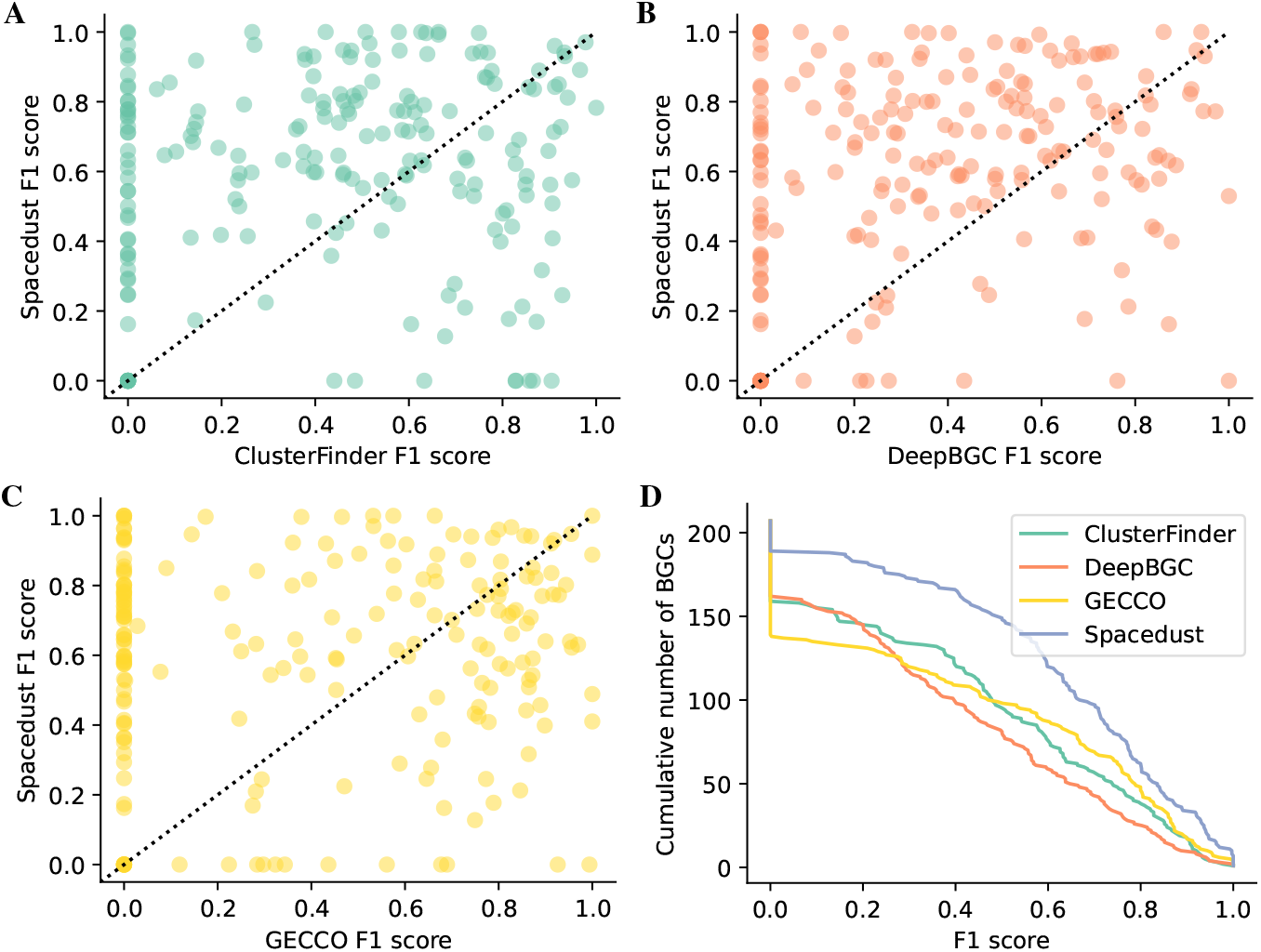
Prediction of 207 manually annotated BGCs from 9 genomes. For each of the 207 biosynthetic gene clusters (BGCs), we compute the F1 score as the harmonic mean of recall and precision. The recall for a BGC is the fraction of genes in the BGC that have been predicted by the tool, and the precision is the fraction of genes in the predicted region that overlap the annotated BGC. (A) Scatter plot of F1 scores of ClusterFinder versus Spacedust, (B) DeepBGC versus Spacedust and (C) GECCO versus Spacedust for the 207 annotated BGCs. (D) Cumulative distribution of the F1 scores for the 207 BGCs.

DeepBGC and GECCO (Fig. 5A-C) due to its higher precision than DeepBGC and GECCO and its higher recall than ClusterFinder (Fig. S8 and S9). All tools failed to detect a few instances of BGCs that were identified by Spacedust. Fig. 5D shows the cumulative distribution of the F1 score for the three tools. The average F1 score over all BGCs is 0.44 for ClusterFinder, 0.39 for DeepBGC, 0.43 for GECCO and 0.61 for Spacedust.

We employed AntiSMASH (28), a tool for profile-based BGC detection, to functionally annotate the genes as either “biosynthetic-related” (biosynthetic, biosynthetic additional, transport, regulatory) or “other genes”. We manually inspected the clusters reported by Spacedust, ClusterFinder, DeepBGC and GECCO. We observed that the regions reported by Spacedust often miss the transport and regulatory genes but cover the core and additional biosynthetic-related genes, sometimes with multiple short gene clusters within the annotated BGC region (Fig. S10).

### Expansion of CRISPR-Cas subtype III-E single effector Cas7-11

Next, we investigated Spacedust’s utility for identifying new instances of known gene cluster families. One such example is the recently discovered CRISPR subtype III-E, which comprises a single effector protein known as Cas7-11 (54). Notably, Cas7-11 is a protein fusing four Cas7 proteins with a putative Cas11-like protein. The fusion yields a single-protein programmable RNase that shows high sequence specificity and no evidence of collateral activity. Previous screening for Cas7-11 across bacterial genomic sequences led to the identification of subtype III-E systems in 17 loci.

To expand our knowledge of subtype III-E systems beyond the reported loci, we queried the proteins in the 17 loci reported by (54) against the GTDB database (55). Because we were unable to map a substantial portion of the query proteins and GTDB proteins to known structures, we employed MMseqs2 iterative search with 3 iterations to perform the homology search. We identified an additional seven instances of subtype III-E clusters in the GTDB database by demanding the presence of the gene cas7-11 (Fig. 6). In three out of seven genomes, all components of the respective system were identified, demonstrating the high sensitivity of the method.

**Fig. 6.**
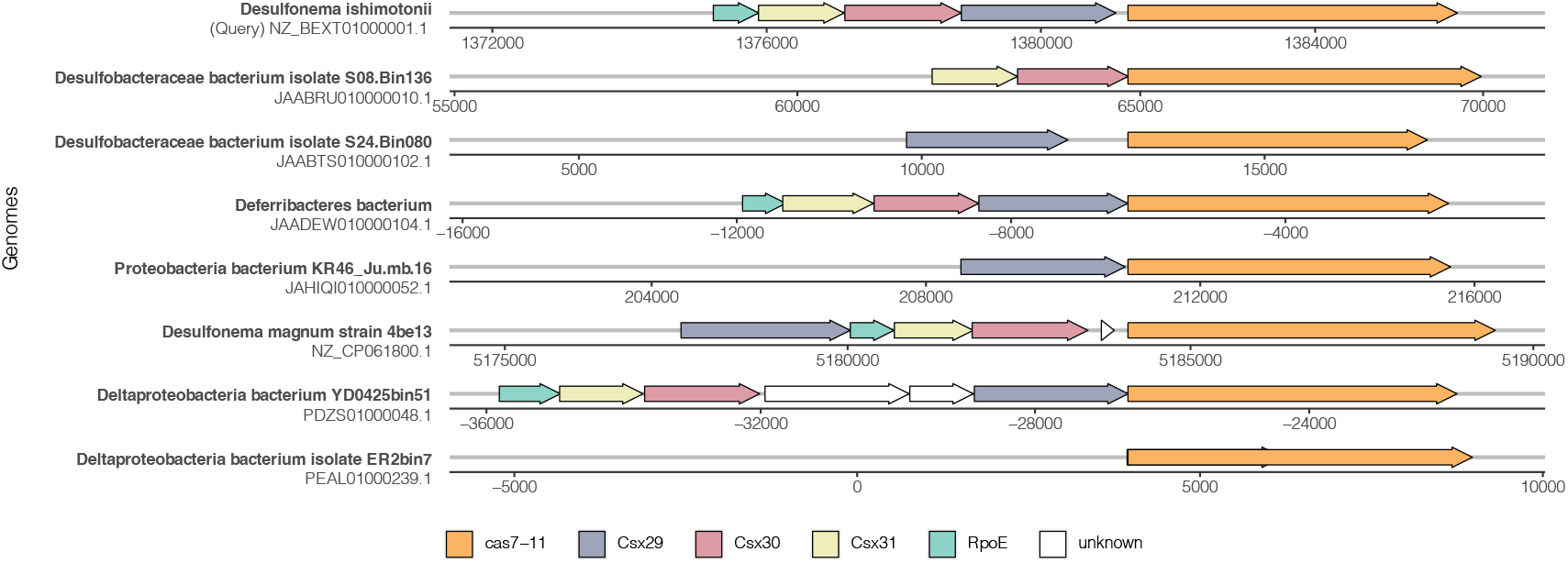
Additional instances of CRISPR-Cas subtype III-E clusters identified in GTDB. Visualization of gene neighborhood of the clusters identified around *cas7-11* (orange arrow). *Desulfonema ishimotonii* (NZ_BEXT01000001.1), the representative of query III-E systems, is plotted as reference to show the gene composition and order. Genes within the cluster boundary that could not be matched to Cas-related genes are colored in white.

### Spacedust Colab Notebook

To facilitate the use of Spacedust for a broad user base, we have set up a Google Colaboratory environment, which allows users to easily run tests and reproduce results without requiring a local installation or configuration. We provide a comprehensive IPython notebook (ipynb file) that includes steps for installing all dependencies and databases, executing the program, and interactively visualizing the clusters. Within the Colab framework, users can easily run Spacedust in either an all-vs-all mode or annotate query genomes against a pre-compiled reference database with just a single click. With the help of interactive visualization, they can explore the evolutionary conservation of their genome of interest in other genomes at different resolutions, and generate gene neighborhood plots for any gene clusters.

## Discussion and Outlook

Exploiting the conservation of gene neighborhoods to predict functional association between genes is an old idea. So far, the main limitations have been (I) the rather low fraction of genes that are part of a conserved gene cluster (44) and (II) the low reliability of the inference of functional association. Spacedust addresses limitation I in three ways. First, it finds homologous proteins using protein structure comparison with Foldseek, which is much more sensitive than the sequence search-based methods used so far. The increased sensitivity yields a higher number of conserved cluster matches between genomes (Figs. S1-3). Second, due to Foldseek’s high speed and Spacedust’s clustered search mode, it can analyze large sets of genomes in an all-versus-all fashion. The large number of genomes increases the chances that a gene will be part of a gene cluster that is conserved in another genome. Certainly, all gene clusters that are laterally transferred as a functional unit – such as biosynthetic gene clusters (56) – will be detectable by Spacedust if a sufficiently large number of genomes is analyzed. Third, Spacedust does not require exact synteny but can find partially conserved neighborhoods (e.g. Fig. 6). The success of these measures is demonstrated by the high fraction of 58 % of genes that are part of a conserved gene cluster among the 4.2 million proteins from the reference genomes of 1 308 bacterial genera (Fig. 2B), as well as the high sensitivities attained for the *de-novo* discovery of antiviral defense systems and BGCs (Figs. 4 and 6).

Spacedust partially addresses limitation II, the low reliability of functional association, by computing two novel P-value statistics for the significance of gene cluster conservation, one assessing the strength of positional clustering of the matched genes and the other assessing the degree of their strand and order conservation. These P-values enable flexibility to find partially conserved gene clusters while still ensuring their statistical significance. Statistically significant conservation alone does not guarantee functional association, however. The fraction of gene pairs within a cluster match that are part of the same KEGG pathway can be as low as 50 % when conserved between only two genomes (Fig 3C). But it rises to over 80 % for the 25 % of the 4.2 million genes that are part of a cluster conserved in at least 50 out of 1 308 genomes (Fig. 2B, C).

The relatively high fraction of KEGG-discordant gene pairs could be due to “false false positives”, functionally associated genes that are not labeled with the same KEGG pathway ID. However, we suspect that the majority of discordant pairs are indeed not functionally associated. This observation was referred to as “genomic hitchhiking” or “carpooling” in (46) and was later rationalized by Fang et al. (19): Hitchhiking, the conservation of genomic neighborhoods containing groups of genes without obvious functional links between them, occurs mainly between core (“persistent”) genes as a side effect of keeping functionally associated, accessory (“non-persistent”) genes clustered together. As a consequence, accessory genes are less involved in hitchhiking. Hitchhiking generally limits the reliability of predicting functional association from conservation of gene neighborhoods.

To assess the sensitivity of Spacedust for finding functionally associated clusters of genes, we compared it with PADLOC, a tool specialized for finding antiviral defense systems. PADLOC relies on a hand-built library of ~ 3800 HMMs to search for the proteins forming part of one of the 210 defense systems. Spacedust discovered *de novo* 95 % of the defense system clusters annotated by PADLOC, out of which 93 % in their entirety (Fig. 4). Similarly, when assessing Spacedust on its ability to discover biosynthetic gene clusters (BGCs) manually annotated in 9 genomes, it performed better than GECCO, DeepBGC and ClusterFinder, which are trained on a large dataset of known BGCs (Fig. 5). In summary, Spacedust has similar sensitivity for *de novo* discovery of functional modules as dedicated tools trained for the discovery of a specific type of gene clusters. However, it is important to note that these specialized tools provide additional value by annotating clusters with specific biosynthetic classes or defense system types, which is not feasible with Spacedust.

The following four limitations of Spacedust need to be addressed in future work: First, partial conservation of a gene cluster in a certain number of genomes predicts functional conservation with only moderate precision (Fig. 2C and D). We are working on an improved conservation score that takes the evolutionary divergence times between genomes into account. We also plan to increase precision by integrating operon predictions on all input genomes. Second, while the fast transformer tool, ProstT5, can reliably predict 3Di sequences for well-studied reference bacterial proteins, its accuracy is less consistent for viral and metagenomic sequences. Although we provide the ProstT5 model to enable full Foldseek structure searches as an alternative to mapping precomputed structures, we emphasize this limitation to guide users in selecting appropriate modes based on their dataset. We are actively working to update the model and will ensure users can easily download the improved version once available. Third, Spacedust cannot find protein members of functional modules encoded outside of a conserved gene cluster (26). We will address this limitation by applying Spacedust for building a database of module-specific protein families and profile HMMs with HMM-specific acceptance thresholds (similar to Pfam (57)), which should allow us to identify also positionally isolated members of functional modules with high specificity. Another limitation of Spacedust is its quadratic scaling of run time with the total number of genomes and proteins in the genomes to be analyzed, caused by the quadratic time complexity of the all-versus-all comparison of proteomes and of the cluster detection algorithm.

Positional orthologs, that is, orthologs that also have conserved gene neighborhood, are under much stronger evolutionary constraints than orthologs without gene neighborhood conservation (58). We therefore expect functional module-specific protein families to show high functional conservation and to become highly useful for automatic functional annotation using a comprehensive profile HMM database of such modules. We also plan to use Spacedust for the systematic discovery of Über operons or extended gene neighborhoods (46, 59, 60), sets of genes that tend to co-occur in each other’s neighborhood more often than by chance and that tend to participate in the same or related processes in the cell.

In conclusion, Spacedust is an extremely sensitive and fast tool for finding conserved gene clusters in large numbers of genomes. It can be employed for the large-scale discovery of modules of functionally associated genes in prokaryotic and viral genomes and metagenome-assembled genomes (MAGs), as demonstrated here with various examples. Its *de novo* approach and high sensitivity make it particularly interesting for the discovery of novel types of functional modules. It can visualize conserved clusters in a query genome across hundreds of target genomes (Fig. 3A). Its tabular output facilitates its integration into current genome annotation pipelines. It can thereby accelerate the elucidation of the functional capabilities of the millions of prokaryotes and viruses that live in and on our bodies and populate all natural environments.

## Materials and Methods

### Spacedust Workflow

#### Input

Spacedust accepts genomic sequences as multiple FASTA files each containing a single prokaryotic genome or metagenome-assembled genome (MAG). Users can either predict the protein coding sequences from input genome using Prodigal v2.6.3 (61) or provide the corresponding GFF3 annotation files of protein coding regions. Contigs belonging to the same assembly should be contained in a single FASTA file. All protein coding regions are extracted and translated. For each protein sequence, the location index, strand and nucleotide coordinates are stored. For all-vs-all comparisons, only one set of genomes is required. For query-to-reference comparisons, a custom database can be built analogously, or a pre-compiled reference database can be automatically downloaded by Spacedust.

#### Mapping to structure database

To enable structure comparisons, query protein sequences are mapped to the reference structure database provided by Foldseek (48). Currently, Fold-seek supports several structure databases such as AlphaFold DB (UniProt, Proteome, Swiss-Prot), PDB and ESMAtlas30. For each protein, in addition to the amino acid sequence, the structure information, including the 3Di sequence and the *C*_*α*_ coordinates, are stored in the Foldseek database. Each query protein sequence is searched against the amino acid sequences of the structure database using MMseqs2 (50) with a stringent sequence identity cutoff of 0.9 and sequence coverage cutoff of 0.9 (`--min-seq-id 0.9 -c 0.9`), and the respective 3Di sequence and the *C*_*α*_ coordinates of the best match are retained to build a Foldseek-compatible database. Alternatively, Foldseek supports translation between protein sequences and 3Di sequences using the ProstT5 protein language model (62), with which a Foldseek-compatible database can be created for all query protein sequences.

#### Homology search

Spacedust conducts a sensitive search of all query proteins against target proteins using Foldseek and/or MMseqs2. If a Foldseek-compatible database is available, the mapped or translated structures sequences will be used in a Foldseek search, while any remaining unmapped sequences are searched using MMseqs2. The E-value cutoff is set at 0.001, and the query sequence coverage is set at 0.8 (`-e 0.001 -c 0.8 --cov-mode 2`). The results from both searches are merged. Users can also opt for an iterative profile (PSI-BLAST like) search by specifying the number of iterations.

#### Clustered search

Searching against a large sequence or structure database is a time and memory demanding task. To improve the search speed while maintaining high sensitivity, we implemented a clustered search workflow similar to the strategy used in ColabFold (63). The query sequences are searched against the consensus sequence or structure of the clustered version of the reference database. For each query hit to a consensus sequence, we realign the query to its respective cluster members and expand the search results. An additional advantage is the higher sensitivity attained using cluster consensus sequences. The clustered search approach results in a 4-fold speed-up, because only 1 million cluster consensus structures are searched instead of 3.8 million structures from the Bacterial Reference database.

#### Hit filtering and grouping

Each protein in each of the genomes in the query set is searched through the proteins in each of the target genomes *T* with *N*_*T*_ proteins. For each query protein, the hit in *t* with the lowest P-value *p* is identified. If *p* ≤ *p*_0_ (default value 10^*−*7^), we compute the probability that in a comparison with *N*_*T*_ non-homologous proteins no P-value will be below *p* is given by the first-order P-value statistics, 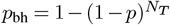. The best hits between the pairs of query and target genomes are then grouped for subsequent cluster detection.

#### Cluster detection

Spacedust employs a probabilistic approach to identify for each pair of genomes conserved neighborhoods of genes, which are clusters of homologous hits with partially conserved clustering and ordering. We assess the conservation of gene neighborhood by combining two P-value statistics, a clustering P-value and an ordering P-value. Given any cluster of hits 𝒞, we can compute the number of hits *k*. The span *m* is the maximum of the number of genes (including unmatched ones) in the query genome cluster and in the target genome cluster. We also define *q*_0_ (default value 10^*−*3^) as the probability for an arbitrary protein *q* to hit an arbitrary protein *t* in *T*. The clustering P-value of the given cluster is the probability to observe a cluster of size *k* each with probability of *q*_0_ within a square of span *m* (Supplementary Material).

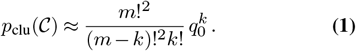

The ordering P-value assesses the statistical significance of the conservation of order and strandedness between two cluster matches. We define the directionality and ordering statistic *n* ∈ {0, 1, 2, …} as the number of neighboring query protein pairs who are also direct neighbors and whose relative orientation is conserved. The ordering P-value is the probability to observe in a randomly occurring cluster match with *k* matched proteins at least *n* neighboring pairs with conserved order and strandedness. In Supplementary Materials section IIB we show that this P-value is

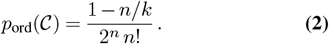

Both statistics are independent of each other and of the strength of individual pairwise sequence homology and thus should improve the specificity of the search. Additionally, both statistics are not influenced by the size of the query and target set, making them suitable for fragmented contigs with small numbers of genes.

The clusters are detected with a greedy agglomerative hierarchical clustering algorithm. Since the two P-values are independent random variables under the null model, we can combine them using the product of P-values *p* := *p*_clu_(𝒞) *p*_ord_(𝒞) (64) which yields the cluster match P-value, *p* × (1 − log *p*). We define a cluster score *S*(𝒞) as negative logarithm of the cluster match P-value: *S*(𝒞) = − log *p*_clu_(𝒞) − log *p*_ord_(𝒞) + log (1 − log *p*_clu_(𝒞) − log *p*_ord_(𝒞). The greedy agglomerative hierarchical clustering algorithm first treats each hit as a singleton cluster, and iteratively merges hits with highest cluster match score satisfying the clustering criteria. Users can adjust the stringency by defining different clustering criteria, such as the maximum number of gaps (non-cluster genes) allowed and the minimum number of genes in a cluster. The probabilistic nature of the algorithm account for micro-rearrangements between genomes, gene insertions/losses and misannotated genes.

#### Output

Spacedust outputs a tab-separated text file. Each reported cluster consists of one summary line followed by multiple lines, one line for each pairwise hit. The summary line starts with ‘#’: a unique cluster identifier, query genome accession, target genome accession, cluster match P-value (joint P-value of clustering and ordering), multihit P-value and number of hits in the cluster. Each following line describes an individual member hit of the cluster in MMseqs2 alignment-result-like format with the following columns: query protein accession, target protein accession, best-hit P-value *p*_bh_, sequence identity, pairwise E-value, query protein start, end and length, target protein start, end and length, alignment traceback string.

### Bacterial Reference database

The bacterial reference database was assembled from KEGG GENOME collection (65), which is comprised of 7,167 complete bacterial genomes. The genomes were downloaded from NCBI GenBank in 09/2022. Genomes were redundancy-filtered using pairwise average amino-acid identities (AAI). Specifically, we used Mash v2.3 (66) to perform all-vs-all alignments using the amino acid alphabet (−a) with default parameters and computed the pairwise AAI as (1 − Mash distance). Next, genomes with AAI of at least 70% were clustered using the SciPy’s hierarchical clustering function (67, 68) and the longest sequence within each cluster was selected as the representative. The threshold roughly corresponds to genus level clustering, meaning the representatives belong to different bacterial genera (69). This resulted in 1 308 representative genomes that make up the reference database. We predicted the protein sequences using Prodigal v2.6.3 and constructed the Foldseek-compatible database as described above. 90.9% (3.8 million of 4.2 million) of the protein sequences could be mapped to a structure in AlphaFold DB. Comprehensive functional annotation of all protein sequences is included using eggNOG-mapper v2.0 (51) with MMseqs2 search and default parameters.

### Reducing the size of reference database

For large reference databases, the search step’s memory requirements would make local runs infeasible. The reference genomes, even after the AAI-based redundancy filtering step, still contain sequence redundancy. Therefore, we further reduce the size of the database by clustering the protein and structure sequences. We clustered the protein sequences with MMseqs2 at 70% sequence identity and 80% bidirectional coverage (`--min-seq-id 0.70 -c 0.8 --cov-mode 0`). For structures, we first clustered the protein sequences at 30% sequence identity and 90% bidirectional coverage with MM-seqs2 (`--min-seq-id 0.30 -c 0.8 --cov-mode 0`), and then further the structure sequences with Foldseek without sequence identity threshold but 90% bidirectional coverage and E-value of less than 0.01. Spacedust provides a search mode `--profile-cluster-search`. Under this search mode, Spacedust only performs MMseqs2 and Foldseek searches against the cluster consensus sequences and then expands to other members of the cluster to not lose hits.

## Data Availability

Spacedust is implemented in C++ and is available as an open-source (GPLv3), user-friendly command-line software for Linux and macOS. The Spacedust source code, compilation instructions, and a user guide are available at https://github.com/soedinglab/Spacedust. Data used in this work was obtained from the public domain and is specified in the respective sections. The bacterial reference genomes were downloaded from NCBI GenBank in 09/2022. The dataset of the 1307 representative bacterial genomes, and scripts to reproduce the search and visualize results are available at https://wwwuser.gwdg.de/~compbiol/spacedust/.

## Acknowledgements

We thank Martin Steinegger (SNU Korea) for his suggestion to use Foldseek in Spacedust. We thank Hong Su (MPI) and Eli Levy Karin for their valuable feedback on the implementation and insightful comments on the manuscript. We used the scientific compute cluster at GWDG, the joint data center of the Max Planck Society (MPG) and University of Göttingen.

## Contributions

RZ and JS designed the Spacedust algorithm, benchmarks, and biological applications. RZ and MM developed the software. RZ performed benchmarks and generated figures. RZ and JS wrote the manuscript. All authors read and approved the final manuscript.

