## Supplementary information for "De novo discovery of conserved gene clusters in microbial genomes with Spacedust"

### Supplementary Material for Spacedust: de novo discovery of conserved gene clusters in microbial genomes

Zhang R.,[1] Mirdita M.,[1,2] and Söding J.[1,3]

[1]*Quantitative and Computational Biology,*
*Max Planck Institute for Multidisciplinary Sciences, Göttingen, Germany*
[2]*School of Biological Sciences, Seoul National University, Seoul, South Korea*
[3]*Campus-Institut Data Science (CIDAS), Göttingen, Germany.*

#### I. SUPPLEMENTARY NOTES

##### A. Eukaryotic genomes

Statistically significant gene clustering has been observed primarily in metabolic gene clusters in fungi and plants [1], and it is reported in multiple other taxa including humans [2]. Spacedust currently does not support input extraction of gene sequences with exon/intron structure from genomic sequences. However, as eukaryotes generally lack operons, these clusters possess different properties in organization and conservation patterns, such as overall much weaker clustering and clustering of unrelated functional categories [3, 4]. Therefore, parameters need to be carefully chosen according to prior knowledge. By leveraging its ability to detect remote homology, Spacedust could contribute to systematically identifying gene clusters conserved across different kingdoms [1]. This is future work.

#### II. SUPPLEMENTARY METHODS

##### A. Cluster definition

In Spacedust, a conserved gene cluster is defined as a group of neighboring genes on a query genome that has matched proteins that are positionally clustered in the target genome, with at least partial conservation of gene neighborhood. Here, the member genes of the cluster need not be strictly in the same order and direction in another genome, and a user-defined number of gaps (non-cluster genes) are allowed between any pairs of members.

#### B. Cluster detection (step 5 in Fig. 1)

Suppose we obtain a list of $K$ pairwise best hits between a query proteome of size $N_Q$ and a target proteome of size $N_T$. So for each $l \in \{1, \ldots, K\}$, we know the position $i_l \in \{1, \ldots, N_\mathrm{Q}\}$ of the query sequence in the query genome and the position $j_l \in \{1, \ldots, N_\mathrm{T}\}$ of the matched target sequence in the target genome.

Our goal is to find clusters of hits $\mathcal{C}$ from $K$ hits such that the hits cluster closely together and have a partially conserved strand and order in both query and target genome. In these cases, the clusters are likely to be part of two groups of functionally associated, positionally orthologous proteins. We assess the degree of conservation of gene neighborhood of any cluster ($\mathcal{C}$) based on two P-value statistics, a clustering P-value $p_{clu}(\mathcal{C})$ and an ordering P-value $p_{ord}(\mathcal{C})$. Both statistics are independent of each other and of individual pairwise sequence similarity, and thus should improve the search sensitivity.

##### *1. Clustering P-value*

We want to assess the degree of clustering of the hits by computing a clustering P-value $p_\mathrm{clu}(\mathcal{C})$. Given any cluster of hits $\mathcal{C} \subset \{1, \ldots, K\}$ that contains $k = |\mathcal{C}|$ hits, we can define its span $m$ as the maximum distance of the cluster members measured as the difference of gene position index on query and target genome:

$$m = \mathrm{span}(\mathcal{C}) = \max\{i_\mathrm{max}(\mathcal{C}) - i_\mathrm{min}(\mathcal{C}), j_\mathrm{max}(\mathcal{C}) - j_\mathrm{min}(\mathcal{C})\}\,. \qquad (1)$$

where

$$\begin{aligned} i_\mathrm{min}(\mathcal{C}) &= \min\{i_l : l \in \mathcal{C}\}, \\ i_\mathrm{max}(\mathcal{C}) &= \max\{i_l : l \in \mathcal{C}\}, \\ j_\mathrm{min}(\mathcal{C}) &= \min\{j_l : l \in \mathcal{C}\}, \\ j_\mathrm{max}(\mathcal{C}) &= \max\{j_l : l \in \mathcal{C}\}. \end{aligned} \qquad (2)$$

The P-value for this statistic is the probability of observing at least $k$ hits in a square of positions with a span of $m$, where each position has a probability of $q_0$ (default $10^{-3}$) of being hit. There are $\binom{m}{k}$ configurations to select $k$ matched proteins from among the $m$ query proteins and the same number to select $k$ matched target proteins from among $m$. There are $k!$ ways to match the $k$ queries to the $k$ targets. The clustering P-value is the probability for a square of $m \times m$ positions in the match matrix to contain $k$ or more matches, where the probability for a cell in this matrix to contain a match is $q_0 \ll 1$. Therefore, the P-value is

$$p_{\text{clu}}(\mathcal{C}) = \sum_{k'=k}^{m} \binom{m}{k'}^2 k'!\, q_0^{k'} = \sum_{k'=k}^{m} \frac{m!^2}{(m-k')!^2 k'!} q_0^k \approx \frac{m!^2}{(m-k)!^2 k!} q_0^k. \tag{3}$$

In the last step, we have approximated the sum with its first term since the second term is smaller than the first by a factor $q_0(m-k)^2/(k+1) \ll 1$, and similarly all other terms are negligible in comparison to the previous one.

The clustering P-value $p_{\text{clu}}(\mathcal{C})$ is independent of the size of query $N_Q$ and target genome $N_T$, which can be applied to fragmented contigs containing few genes. The user can adjust the value of $q_0$ for the stringency of the clustering.

*2. Ordering P-value.*

For clusters of hits $\mathcal{C} \subset \{1, \ldots, K\}$ we want to assess the degree of conservation of their order and transcription directionality (strand) by computing an ordering P-value $p_{\text{ord}}(\mathcal{C})$. For any given cluster $\mathcal{C}$ of $k$ hits, we re-index the hits as $(i_l, j_l), l \in \{1, \ldots, k\}$, with position index in the query set $i_l$ sorted in ascending order, $i_1 < i_2 < \ldots < i_k$. We also denote the direction of transcription (strand of the encoding ORF) as $d_l \in \{-1, +1\}$ (+1 for forward strand, -1 for reverse strand) for the $l$'th query protein and $d'_l \in \{-1, +1\}$ for the corresponding, matched target protein.

We define the directionality and ordering statistic as the number of query protein pairs $(i_l, i_{l+1})$ whose matched target proteins $(j_l, j_{l+1})$ have the same direction and order.

$$n = \sum_{l=1}^{k-1} I\left(d_l\, d'_l = \text{sign}(j_{l+1} - j_l)\right)\, I\left(d_{l+1}\, d'_{l+1} = \text{sign}(j_{l+1} - j_l)\right)\,, \tag{4}$$

where $I(\cdot)$ is the indicator function. The expression in the summation is equal to 1 only in two cases: (1) the target protein (gene) pair have the same order as in the query, $j_{l+1} - j_l > 0$, and the direction of each hit are the same, $d_{l+1}$ and $d'_{l+1}$ have the same sign and $d_l$ and $d'_l$ have the same sign, (2) the target protein(gene) pair have the reversed order as in the query,

$j_{l+1} - j_l < 0$, and $d_l$ and $d'_l$ have opposite signs. The second case occurs when the genes are encoded on opposite strands in the query and target genomes, i.e., $(+,-)$ or $(-,+)$.

The ordering P-value is equal to the number of configurations under $n$ constraints for $k-1$ pairs divided by the total number of possible configurations. For a cluster of $k$ hits, there are $2^k\,k!$ total possible combinations of ordering and direction. But since the choice of forward strand in the target genome is arbitrary, we only need to consider half of the configurations, $C(k) = 2^{k-1}\,k!$. For a conserved pair of hits $(i_l, i_{l+1})$ the constraint determines the sign of $j_{l+1} - j_l$ and of $d_{l+1}d'_{l+1}$ given the sign of $d_l d'_l$. Therefore, each pair with such a constraint behaves in a combinatorial sense as a single transcription unit, and therefore each such constraint reduces the effective number of independent transcription units by 1. According to equation (4) the number of these constraints is $n$. There are $k-1$ pairs which could be constrained in such a way, hence there are $\binom{k-1}{n}$ configurations in which at least $n$ pairs are constrained. So the number of configurations under $n$ or more constraints is therefore $C(k-n)\binom{k-1}{n}$.

The ordering P-value is the number of configurations under at least $n$ constraints $C(k-n)\binom{k-1}{n}$ divided by the total number $C(k) = 2^{k-1}\,k!$ of possible configurations :

$$p_{\text{ord}}(\mathcal{C}) = \frac{C(k-n)}{C(k)}\binom{k-1}{n} = \frac{2^{k-n-1}(k-n)!}{2^{k-1}\,k!}\frac{(k-1)!}{(k-n-1)!\,n!} \quad (5)$$

or

$$p_{\text{ord}}(\mathcal{C}) = \frac{1-n/k}{2^n\,n!}\,. \quad (6)$$

*3. Multihit P-value*

We consider a cluster composed of $k$ pairwise hit with best-hit P-values $p_1, \ldots, p_k$. In order to provide a summary statistic of the homologous relationship of a cluster, we need to take into account the strength and the number of individual hits. We assume that if a sizable fraction of the hits have a significant similarity, it already constitutes a signal for a global significant similarity. A standard approach is to use the Fisher's method [5] for combining $k$ independent P-values of the individual hits that are the product of all P-values:

$$F = \prod_{i=1}^{k} p_i \quad (7)$$

However, the Fisher's method is not suitable when many P-values in the cluster are not significant, as they will dilute the actual significant homology signals. In order to tackle the issue, the previous works introduced a truncated version of the Fisher's method that takes into account only P-values below a given $p_0$ threshold [7]. As a drawback, the truncated $F$ statistic has a strong discontinuity when a single P-value falls below the threshold. We recall that each or the pairwise hits between sequences in the query and target sets has to have a best-hit P-value less than or equal to some threshold $p_0 = 1/(N_{\mathrm{T}} + 1)$. We consider a cluster composed of $k$ pairwise hit with P-values $p_1, \ldots, p_k < p_0$. We propose a variant $R$ of the truncated $F$ statistic, that allows a continuous increase of the R-value by summing up the parts of evidence that pass the threshold $p_0$:

$$R = -\sum_{i=1}^{k} \log \frac{p_i}{p_0} \tag{8}$$

Because the P-values of pairwise hits are uniformly distributed over $[0, 1]$ under the null model, the subset of P-values smaller than $p_0$ are also uniformly distributed, and the $p_i/p_0$ are therefore uniformly distributed on $[0, 1]$. The multi-hit R statistic can distinguish homology signals from random cases, while still reflecting the strength and number of individual hits. The P-value for a variable equal to the product of these $k$ independent P-values is well known [5, 6]:

$$\begin{aligned} \text{P-value}(e^{-r}) &= p\left(\prod_{i=1}^{k} \frac{p_i}{p_0} \le e^{-r} \middle| p_1, \ldots, p_k \sim U(0, p_0)\right) \\ &= p\left(\prod_{i=1}^{k} p'_i \le e^{-r} \middle| p'_1, \ldots, p'_k \sim U(0, 1)\right) \\ &= e^{-r} \sum_{i=0}^{k-1} \frac{r^i}{i!}\,, \end{aligned} \tag{9}$$

where r follows the R statistic.

This multihit P-value is distributed according to $U(0, 1)$ under the null model. It has some similarity with the standard E-value used for pairwise matches in sequence searching: It already takes account of the size of the target set (by dint of the cut-off $p_0$ on the pairwise hit P-values), and it does not correct for multiple testing with respect to the multiple queries.

##### C. Greedy hierarchical agglomerative clustering algorithm (step 6 in Fig. 1)

This algorithm is used for detecting cluster of hits with conserved gene neighborhood. We obtain the clustering and ordering P-values $p_{clu}(\mathcal{C})$ and $p_{ord}(\mathcal{C})$ separately for any cluster of hits $\mathcal{C}$, of which the calculation is detailed in the subsections II B 1 and II B 2. These two P-values are independent random variables under the null model. We can therefore combine them using the product of P-values[6]

$$p := p_{\text{clu}}(\mathcal{C})\, p_{\text{ord}}(\mathcal{C}). \tag{10}$$

The P-value of $p$, termed cluster match P-value, is

$$f(p) = p(1 - \log p). \tag{11}$$

We can define a cluster score $S(\mathcal{C})$ as the minus natural logarithm of the cluster match P-value $f(p)$:

$$S(\mathcal{C}) := -\log f(p) = -\log p_{\text{clu}}(\mathcal{C}) - \log p_{\text{ord}}(\mathcal{C}) + \log\left(1 - \log p_{\text{clu}}(\mathcal{C}) - \log p_{\text{ord}}(\mathcal{C})\right). \tag{12}$$

We propose a greedy algorithm for identifying cluster $\mathcal{C}$ of pairwise hits with a maximum score $S(\mathcal{C})$, since finding the cluster of minimum P-value is equivalent to maximizing the score $S(\mathcal{C})$. Due to the monotonicity of $f(p)$, we only need to maximize the first part of the equation $-\log p_{\text{clu}}(\mathcal{C}) - \log p_{\text{ord}}(\mathcal{C})$, which is faster to compute in practice.

The agglomerative hierarchical clustering algorithms treat each hit as a singleton cluster in the beginning and then successively merge pairs of clusters until all clusters have been merged into a single cluster that contains all hits [8]. The algorithm proceeds as follows:

1. Add each hit $l$ as a singleton cluster to the list of clusters $\mathcal{L}$.

2. Compute the scores $S(\mathcal{C} \cup \mathcal{C}')$ using equation (12) for all pairs of *compatible* clusters $\mathcal{C}, \mathcal{C}' \in \mathcal{L}$, where compatibility requires that at most $d-1$ genes lie between the genes or the two clusters in the query genome and likewise in the target genome,

$$\begin{aligned} &\min\{|j_{\min}(\mathcal{C}) - j_{\max}(\mathcal{C}')|\,, |j_{\min}(\mathcal{C}') - j_{\max}(\mathcal{C})|\} \leq d \text{ and} \\ &\min\{|i_{\min}(\mathcal{C}) - i_{\max}(\mathcal{C}')|\,, |i_{\min}(\mathcal{C}') - i_{\max}(\mathcal{C})|\} \leq d \end{aligned} \tag{13}$$

and no protein in the query set or target set must be matched more than once, i.e $\forall\, l \in \mathcal{C}, l' \in \mathcal{C}' : i_l \neq i_{l'} \wedge j_l \neq j_{l'}$.

3. While the largest score $S(\mathcal{C} \cup \mathcal{C}')$ for merging compatible clusters exceeds some merging threshold $S_{\text{merge}}$,

4. remove $\mathcal{C}$ and $\mathcal{C}'$ from $\mathcal{L}$ and all their merge scores with other clusters from the score list;

5. add $\mathcal{C} \cup \mathcal{C}'$ to $\mathcal{L}$ and add the merge scores of $\mathcal{C} \cup \mathcal{C}'$ with all compatible clusters in $\mathcal{L}$ to the score list.

6. Return all non-singleton clusters in $\mathcal{L}$ whose score is above a significance threshold $S_{\text{min}}$.

The merging threshold $S_{merge}$ is derived from $d$ as the maximum cluster score of a cluster of two elements with a span of $d - 1$. The maximum number of genes allowed between two clusters $d$ and the significance threshold $S_{min}$ for reporting the clusters are both user-definable parameters, where $S_{min}$ is converted to cluster match P-value as in equation (11). Simultaneously, the reported cluster has to satisfy a user-defined multihit P-value threshold, of which the computation is detailed in subsection II B 3.

##### *1. Gene fusion*

In the agglomerate clustering step, we do not allow any query or target protein to be matched more than once. Comparative genomic revealed cases in which two or more proteins encoded separately in one genome appear as fusions either in the same genome or some other genomes. Such fusion proteins, termed Rosetta stone sequences, are also strongly suggestive of functional relationships of the disparate proteins, and thus should be detected and reported in the cluster output [9].

On the sequence level, a gene fusion case has to qualify for the following criteria: (1) at least two hits match to the same target protein (2) the hits do not overlap or only overlaps very little on the target protein (3) a significant part of the target protein is covered by the query proteins (4) a significant part of the query proteins are covered by the target protein and (5) the order and direction of transcription is retained. In the homology search step, we demand a query coverage of $\geq$80%. Only one best hit in target set is chosen for each query protein.

To include gene fusion events, we allow two proteins encoded by consecutive genes at position $i_l, i_{l+1}$ in query set to be matched to one protein $j_{l'}$ in the target, only if the two aligned regions in the one protein do not overlap, and the aligned region has to cover $\geq$80%

of the target protein, and the order and direction of transcription are conserved.

#### III. COMPUTATIONAL RESOURCES

All runs are performed on the GWDG HPC cluster (www.gwdg.de) using individual nodes with two Intel Xeon E5-2640v3 processors at 2.6 GHz each possessing $2 \times 64$ cores and 1024 GB RAM.

#### IV. SOFTWARE VERSIONS

| Name | Version |
|---:|---:|
| Spacedust | Git: 2-e56c505 |
| Foldseek | 10-941cd33 |
| ProstT5 | v0.0.1 |
| Prodigal | 2.6.3 |
| eggNOG-mapper | v2.0 |
| Mash | v2.3 |
| PADLOC | v1.1.0 |
| ClusterFinder | Git: 5ee2c15 |
| DeepBGC | v0.1.29 |
| AntiSMASH | v6.0.0 |
| GECCO | v0.9.10 |

TABLE I. Software versions used in this manuscript.

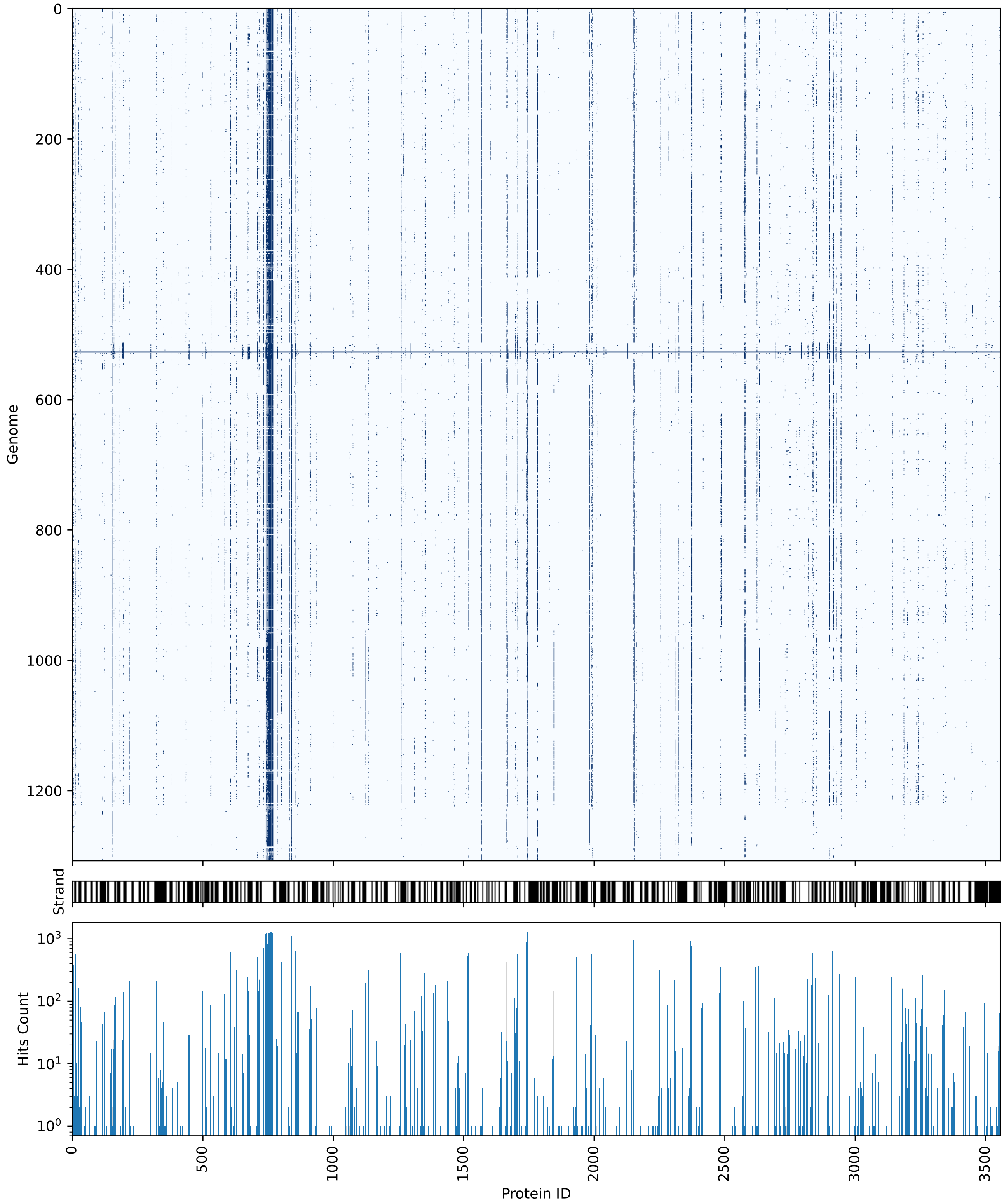


FIG. 1. Evolutionary conservation of gene clusters in a cyanobacterium *Synechocystis* sp. PCC6803. Clustered hits of *Synechocystis* sp. PCC6803 (Genome ID 527) against 1308 bacterial reference genomes using Spacedust Foldseek+MMseqs2 search.

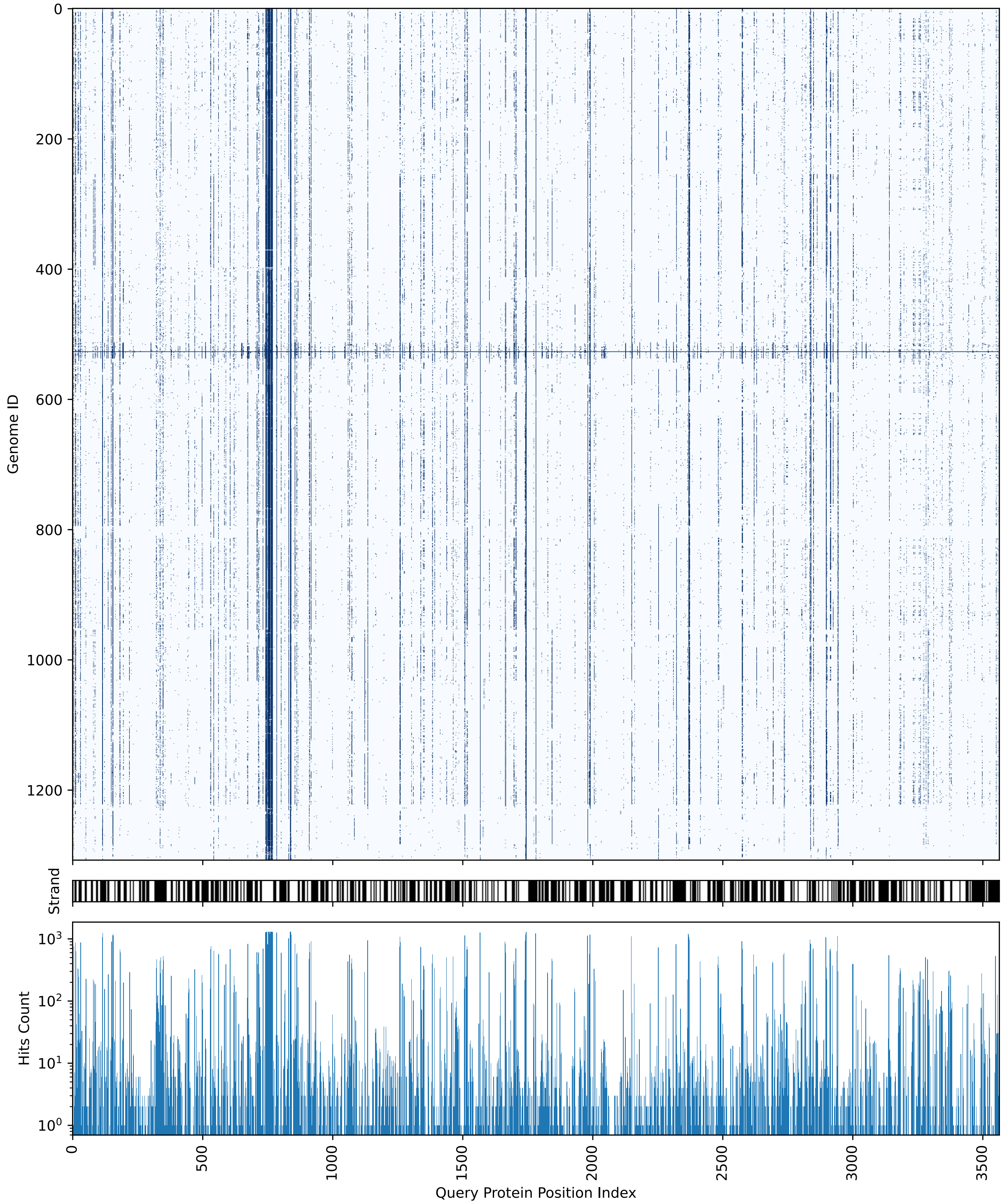


FIG. 2. Evolutionary conservation of gene clusters in a cyanobacterium *Synechocystis* sp. PCC6803. Clustered hits of *Synechocystis* sp. PCC6803 (Genome ID 527) against 1308 bacterial reference genomes using Spacedust Foldseek-only search with 3Di sequences predicted by ProstT5.

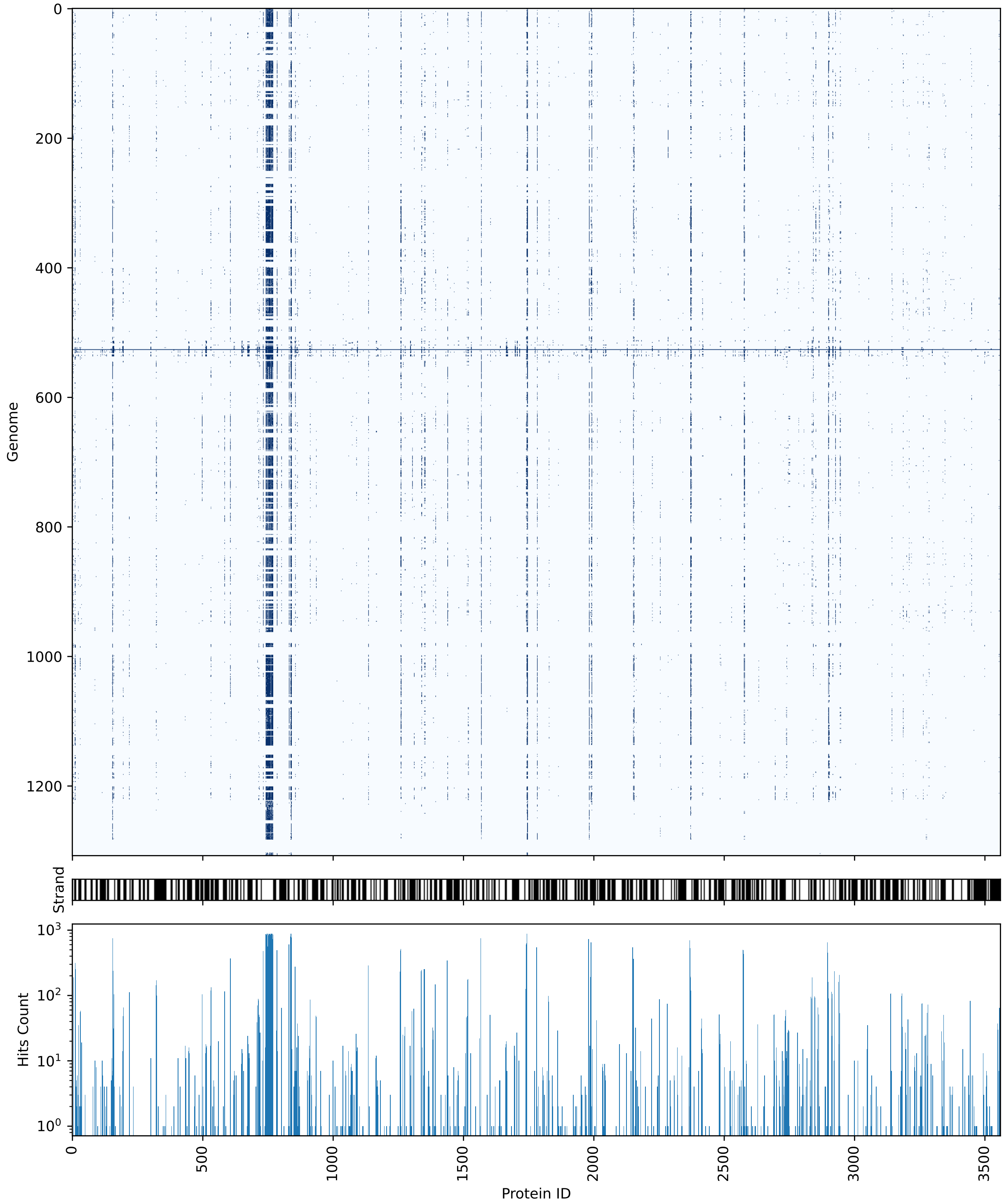


FIG. 3. Evolutionary conservation of gene clusters in a cyanobacterium *Synechocystis* sp. PCC6803. Clustered hits of *Synechocystis* sp. PCC6803 (Genome ID 527) against 1308 bacterial reference genomes using Spacedust MMseqs2 search.

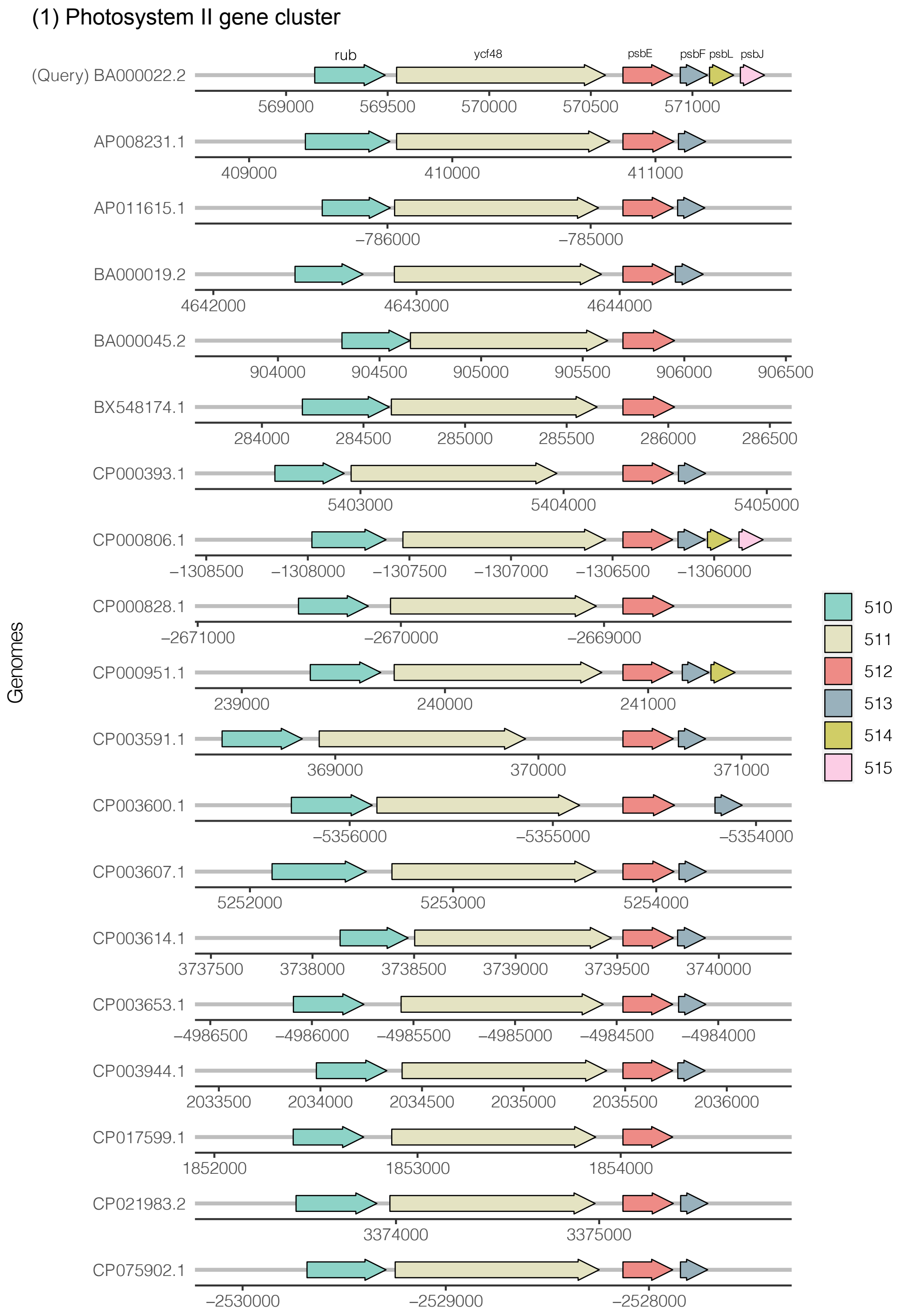


FIG. 4. Gene neighborhood of Cyanobacteria-specific cluster 1 (Protein ID 510-515), centered around protein 512

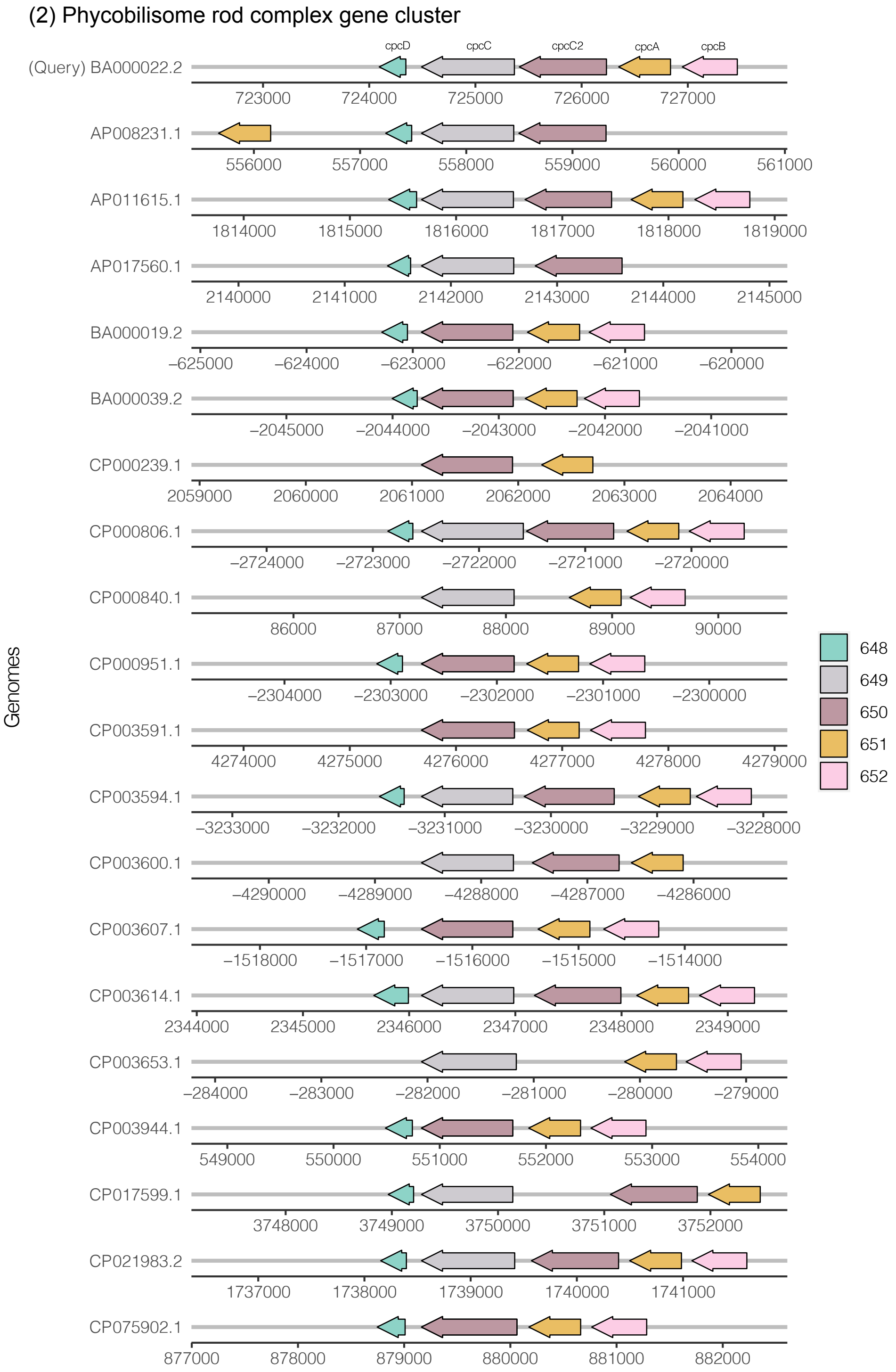


FIG. 5. Gene neighborhood of Cyanobacteria-specific cluster 2 (Protein ID 648-652), centered around protein 649

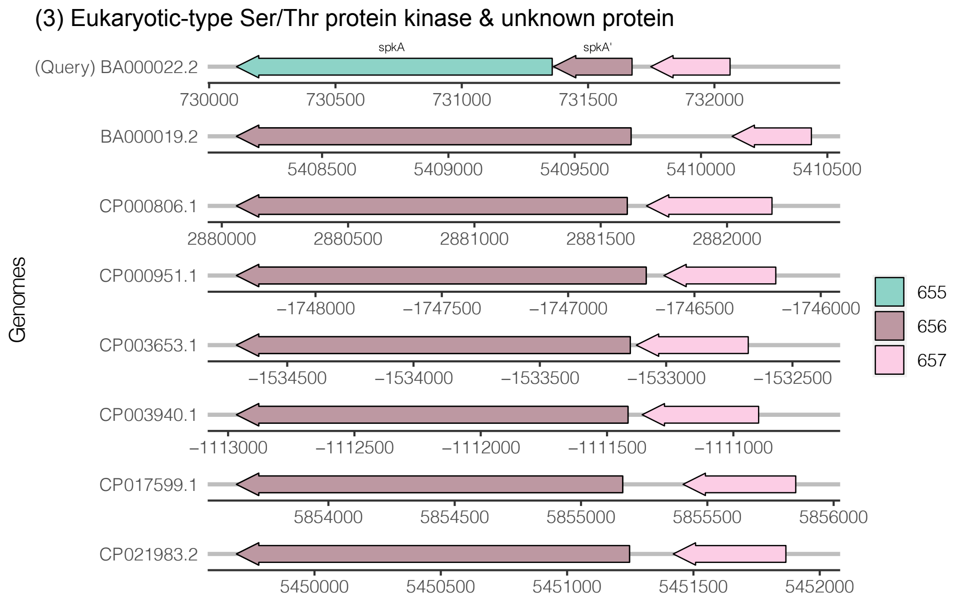


FIG. 6. Gene neighborhood of Cyanobacteria-specific cluster 3 (Protein ID 655-657), centered around protein 655

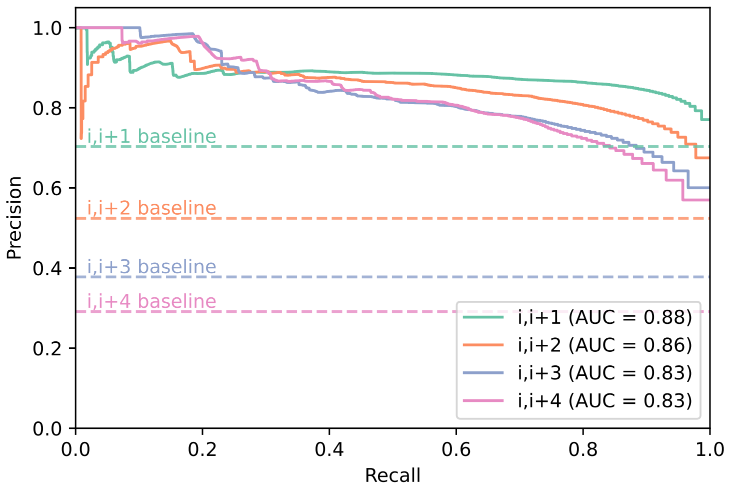


**(A)**

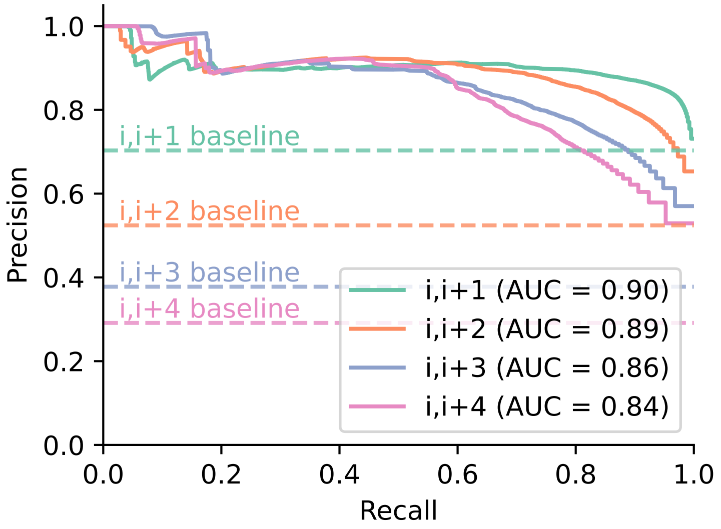


**(B)**

FIG. 7. Precision-recall (PR) of functional association of non-redundant conserved clusters (including the ribosomal genes) for (A) Foldseek+MMseqs search and (B) Foldseek-only search with 3Di sequences predicted by ProstT5, assessed by congruence of KEGG module IDs of Spacedust cluster matches for all gene pairs separated by up to 4 genes (i,i+1),..., (i,i+4).

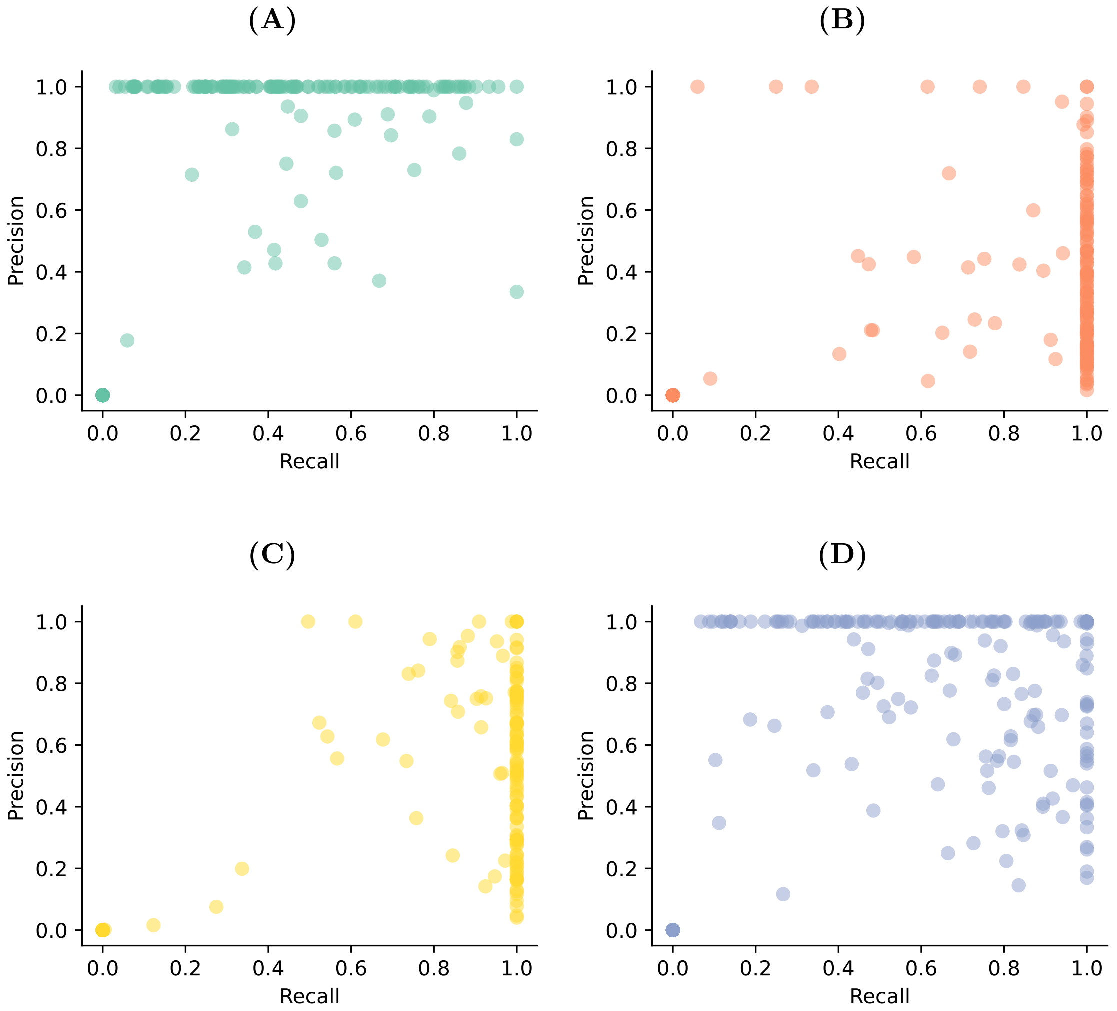


FIG. 8. Scatter plots of precision versus recall for the 207 annotated BGCs for (A) Clusterfinder, (B) DeepBGC, (C) GECCO and (D) Spacedust.

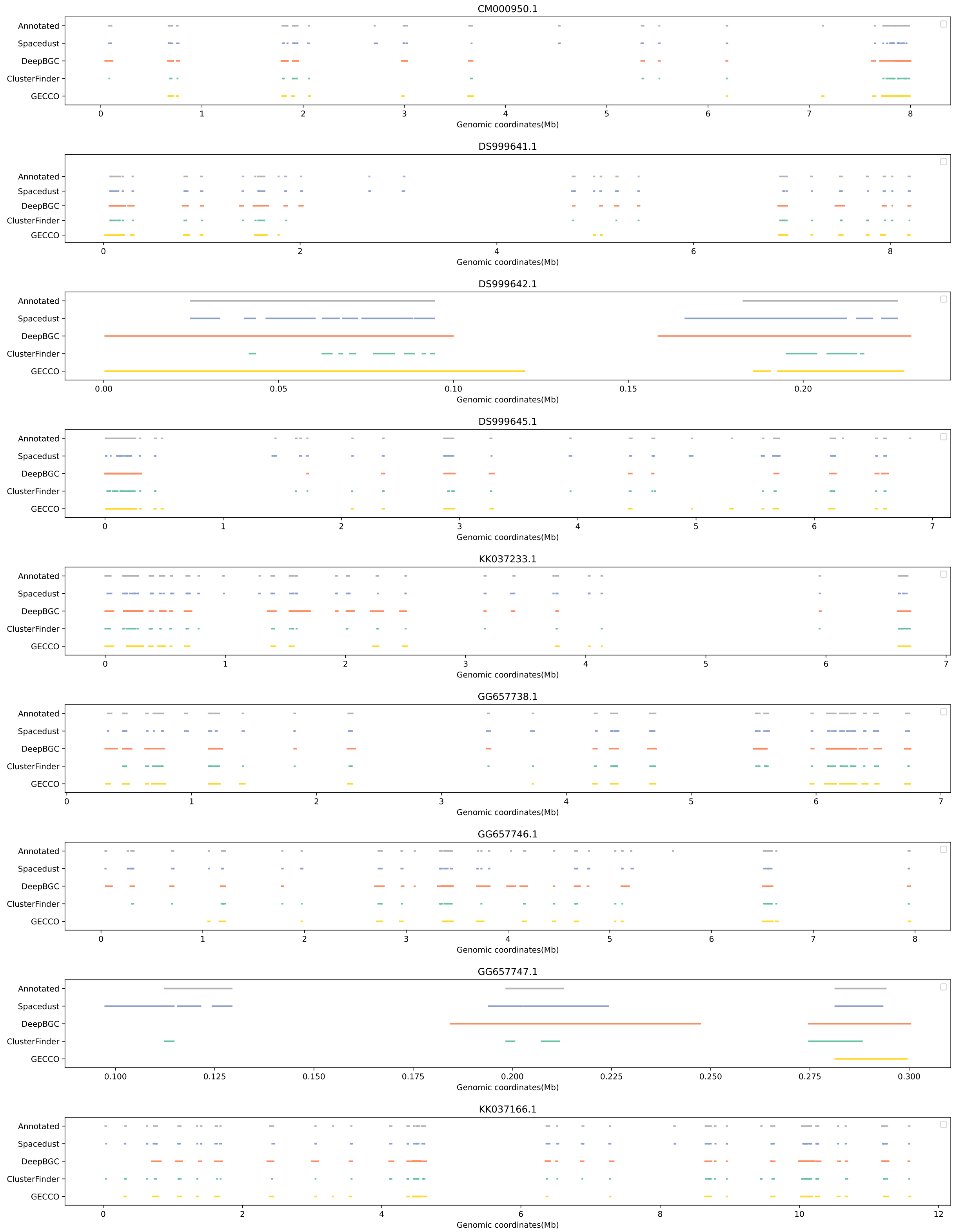


FIG. 9. Contig view of 9 reference genomes with genomic regions predicted by ClusterFinder (green), DeepBGC (orange), GECCO (yellow) and Spacedust (blue) overlapping with the annotated BGCs (grey).

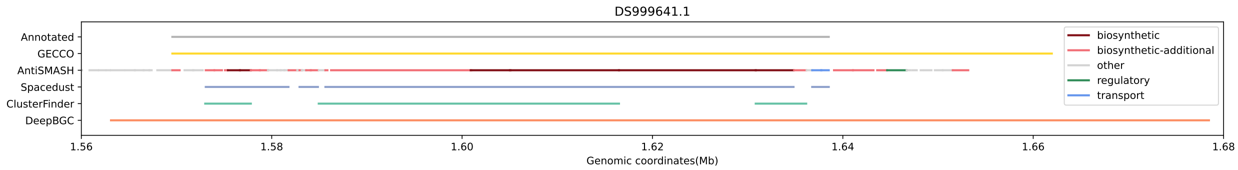


FIG. 10. Example BGC regions (DS999641.1) identified by ClusterFinder (green), DeepBGC (orange), GECCO (Yellow) and Spacedust (blue), superimposed upon annotated BGCs (grey) along with AntiSMASH's predictions and functional categories.

TABLE II. EggNOG-mapper annotation of Synechocystis sp. PCC6803 clusters

| Index | Start | End | COG category | Name | EC | KEGG_ko | KEGG_Module | PFAMs |
|---|---|---|---|---|---|---|---|---|
| 510 | 569141 | 569488 | C | rub | - | - | - | Rubredoxin |
| 511 | 569544 | 570572 | S | ycf48 | - | - | - | PSII_BNR |
| 512 | 570658 | 570903 | C | psbE | - | ko:K02707 | M00161 | Cytochrom_B559,Cytochrom_B559a |
| 513 | 570940 | 571074 | A | psbF | - | ko:K02708 | M00161 | Cytochrom_B559 |
| 514 | 571084 | 571203 | U | psbL | - | ko:K02713 | - | PsbL |
| 515 | 571236 | 571355 | U | psbJ | - | ko:K02711 | - | PsbJ |
| 528 | 583092 | 582481 | S | - | - | - | - | Uma2 |
| 529 | 583573 | 583157 | P | - | - | ko:K03711 | - | FUR |
| 530 | 583680 | 584696 | P | zntC | - | ko:K09815 | M00242 | ZnuA |
| 531 | 584674 | 585543 | P | - | - | ko:K09817 | M00242 | ABC_tran |
| 532 | 585497 | 586342 | U | - | - | ko:K09816 | M00242 | ABC-3 |
| 542 | 600833 | 599910 | J | dnaJ3 | - | ko:K05516 | - | DnaJ,DnaJ_C |
| 543 | 603308 | 600942 | O | dnaK1 | - | ko:K04043 | - | HSP70 |
| 583 | 644379 | 643549 | S | - | 3.1.2.12 | ko:K01070 | - | Esterase |
| 584 | 645497 | 644388 | C | frmA | 1.1.1.1,1.1.1.284 | ko:K00121 | - | ADH_N,ADH_zinc_N |
| 586 | 658262 | 659335 | H | chlI | 6.6.1.1 | ko:K03405 | - | Mg_chelatase |
| 587 | 659370 | 660584 | J | tyrS | 6.1.1.1 | ko:K01866 | M00359,M00360 | S4,tRNA-synt_1b |
| 590 | 662952 | 663188 | J | tyrS | 6.1.1.1 | ko:K01866 | M00359,M00360 | S4,tRNA-synt_1b |
| 591 | 663342 | 663731 | L | ycf41 | - | ko:K03111 | - | SSB |
| 605 | 673889 | 675340 | C | cydA | 1.10.3.14 | ko:K00425 | M00153 | Cyt_bd_oxida_I |
| 606 | 675344 | 676354 | C | cydB | 1.10.3.14 | ko:K00426 | M00153 | Cyt_bd_oxida_II |
| 608 | 678286 | 677045 | F | ackA | 2.7.2.1 | ko:K00925 | M00357,M00579 | Acetate_kinase |
| 609 | 679023 | 678283 | Q | clcD | 3.1.1.45 | ko:K01061 | - | DLH |
| 616 | 684156 | 683140 | P | pobA | - | - | - | Rieske |
| 617 | 684439 | 684143 | C | - | - | - | - | Fer2 |
| 618 | 687334 | 684560 | NT | - | - | ko:K02487 | M00507 | CheW,H-kinase_dim,HATPase_c,Hpt,Response_reg |
| 619 | 690252 | 687391 | NT | pilJ | - | ko:K02660 | - | GAF,HAMP,MCPsignal |
| 620 | 690866 | 690399 | NT | - | - | ko:K02659 | - | CheW |
| 621 | 691264 | 690899 | KT | - | - | ko:K02658 | M00507 | Response_reg |
| 622 | 692576 | 691371 | KT | - | - | ko:K02657 | M00507 | DUF4388,Response_reg |
| 624 | 695874 | 695290 | O | - | - | - | - | AhpC-TSA |
| 627 | 698723 | 698962 | P | - | - | ko:K04758 | - | FeoA |
| 628 | 698970 | 700814 | P | feoB | - | ko:K04759 | - | FeoB_C,FeoB_N,Gate |
| 631 | 704390 | 704845 | S | - | - | ko:K09775 | - | DUF212 |

TABLE III. EggNOG-mapper annotation of Synechocystis sp. PCC6803 clusters (continued)

| Index | Start | End | COG category | Name | EC | KEGG_ko | KEGG_Module | PFAMs |
|---|---|---|---|---|---|---|---|---|
| 648 | 724345 | 724094 | P | cpcD | – | ko:K02287 | – | CpcD |
| 649 | 725367 | 724492 | H | cpcC | – | ko:K02286 | – | CpcD,PBS_linker_poly |
| 650 | 726234 | 725413 | H | cpcC2 | – | ko:K02286 | – | CpcD,PBS_linker_poly |
| 651 | 726837 | 726349 | C | cpcA | – | ko:K02284 | – | Phycobilisome |
| 652 | 727466 | 726948 | C | cpcB | – | ko:K02285 | – | Phycobilisome |
| 655 | 731358 | 730108 | KLT | spkA | 2.7.11.1 | ko:K12132 | – | Pkinase |
| 656 | 731674 | 731363 | KLT | spkA | 2.7.11.1 | ko:K12132 | – | Pkinase |
| 657 | 732062 | 731748 | – | – | – | – | – | – |
| 671 | 747234 | 747578 | P | – | – | ko:K05567 | – | Oxidored_q2 |
| 672 | 747575 | 749038 | CP | ndhD5 | – | ko:K05568 | – | Proton_antipo_M |
| 673 | 749150 | 749500 | P | – | – | ko:K05567 | – | Oxidored_q2 |
| 674 | 749550 | 751037 | CP | ndhD5 | – | ko:K05568 | – | Proton_antipo_M |
| 675 | 751034 | 751426 | P | – | – | ko:K05569 | – | MNHE |
| 676 | 751381 | 751674 | – | – | – | ko:K05570 | – | – |
| 677 | 751671 | 751967 | P | – | – | ko:K05571 | – | PhaG_MnhG_YufB |
| 678 | 751936 | 752535 | P | – | – | ko:K07242 | – | DUF4040 |
| 679 | 752532 | 753179 | P | – | – | ko:K05566 | – | MnhB |
| 682 | 756053 | 757195 | NU | – | – | – | – | N_methyl |
| 683 | 757244 | 757720 | NU | – | – | – | – | N_methyl |
| 684 | 757671 | 758753 | NU | – | – | – | – | N_methyl |
| 685 | 758829 | 761228 | NU | – | – | – | – | – |
| 708 | 790719 | 791717 | C | coxB | 1.9.3.1 | ko:K02275 | M00155 | COX2,COX2_TM |
| 709 | 791806 | 793461 | C | ctaDI | 1.9.3.1 | ko:K02274 | M00155 | COX1 |
| 710 | 793509 | 794210 | C | ctaE | 1.9.3.1 | ko:K02276 | M00155 | COX3 |
| 713 | 796732 | 795932 | P | tauB | – | ko:K02049 | M00188 | ABC_tran |
| 714 | 797631 | 796789 | P | – | – | ko:K02050 | M00188 | BPD_transp_1 |
| 715 | 798792 | 797734 | P | – | – | ko:K02051 | M00188 | NMT1,NMT1_2 |
| 716 | 799535 | 798846 | KO | hypB | – | ko:K04652 | – | cobW |
| 717 | 799958 | 799596 | S | hypA | – | ko:K04651 | – | HypA |
| 718 | 801206 | 800034 | E | speB2 | 3.5.3.11 | ko:K01480 | M00133 | Arginase |
| 719 | 801643 | 802152 | T | – | – | – | – | GAF |
| 720 | 802207 | 803238 | T | – | – | – | – | GAF,GGDEF |
| 730 | 819979 | 817967 | G | tktA | 2.2.1.1 | ko:K00615 | M00004,M00007,M00165,M00167 | Transket_pyr,Transketolase_C,Transketolase_N |
| 731 | 821352 | 820102 | I | fabF | 2.3.1.179 | ko:K09458 | M00083,M00572 | Ketoacyl-synt_C,ketoacyl-synt |
| 732 | 821783 | 821550 | IQ | acpP | – | ko:K02078 | – | PP-binding |

TABLE IV. EggNOG-mapper annotation of Synechocystis sp. PCC6803 clusters (continued)

| Index | Start | End | COG category | Name | EC | KEGG_ko | KEGG_Module | PFAMs |
|---|---|---|---|---|---|---|---|---|
| 740 | 826882 | 826637 | J | rpmE | - | ko:K02909 | M00178 | Ribosomal_L31 |
| 741 | 827348 | 826935 | J | rps9 | - | ko:K02996 | M00178,M00179 | Ribosomal_S9 |
| 742 | 827803 | 827348 | J | rplM | - | ko:K02871 | M00178,M00179 | Ribosomal_L13 |
| 743 | 828695 | 827868 | J | truA | 5.4.99.12 | ko:K06173 | - | PseudoU_synth_1 |
| 744 | 829063 | 828713 | J | rplQ | - | ko:K02879 | M00178 | Ribosomal_L17 |
| 745 | 830085 | 829141 | K | rpoA | 2.7.7.6 | ko:K03040 | M00183 | RNA_pol_A_CTD,RNA_pol_A_bac,RNA_pol_L |
| 746 | 830586 | 830194 | J | rpsK | - | ko:K02948 | M00178,M00179 | Ribosomal_S11 |
| 747 | 831018 | 830635 | J | rpsM | - | ko:K02952 | M00178,M00179 | Ribosomal_S13 |
| 748 | 831217 | 831101 | J | rpmJ | - | ko:K02919 | M00178 | Ribosomal_L36 |
| 749 | 831558 | 831355 | J | infA | - | ko:K02518 | - | eIF-1a |
| 750 | 832265 | 831702 | F | adk | 2.7.4.3 | ko:K00939 | M00049 | ADK |
| 751 | 833719 | 832391 | U | secY | - | ko:K03076 | M00335 | SecY |
| 752 | 834264 | 833821 | J | rplO | - | ko:K02876 | M00178,M00179 | Ribosomal_L27A |
| 753 | 834854 | 834333 | J | rps5 | - | ko:K02988 | M00178,M00179 | Ribosomal_S5,Ribosomal_S5_C |
| 754 | 835256 | 834894 | J | rplR | - | ko:K02881 | M00178,M00179 | Ribosomal_L18p |
| 755 | 835799 | 835260 | J | rpl6 | - | ko:K02933 | M00178,M00179 | Ribosomal_L6 |
| 756 | 836260 | 835859 | J | rps8 | - | ko:K02994 | M00178,M00179 | Ribosomal_S8 |
| 757 | 836954 | 836352 | J | rpl5 | - | ko:K02931 | M00178,M00179 | Ribosomal_L5,Ribosomal_L5_C |
| 758 | 837348 | 837001 | J | rplX | - | ko:K02895 | M00178,M00179 | KOW,ribosomal_L24 |
| 759 | 837717 | 837349 | J | rplN | - | ko:K02874 | M00178,M00179 | Ribosomal_L14 |
| 760 | 837986 | 837741 | J | rpsQ | - | ko:K02961 | M00178,M00179 | Ribosomal_S17 |
| 761 | 838215 | 837994 | J | rpmC | - | ko:K02904 | M00178,M00179 | Ribosomal_L29 |
| 762 | 838637 | 838218 | J | rplP | - | ko:K02878 | M00178 | Ribosomal_L16 |
| 763 | 839400 | 838678 | J | rps3 | - | ko:K02982 | M00178,M00179 | KH_2,Ribosomal_S3_C |
| 764 | 839795 | 839430 | J | rplV | - | ko:K02890 | M00178,M00179 | Ribosomal_L22 |
| 765 | 840089 | 839811 | J | rpsS | - | ko:K02965 | M00178,M00179 | Ribosomal_S19 |
| 766 | 840952 | 840122 | J | rpl2 | - | ko:K02886 | M00178,M00179 | Ribosomal_L2,Ribosomal_L2_C |
| 767 | 841307 | 841002 | J | rplW | - | ko:K02892 | M00178,M00179 | Ribosomal_L23 |
| 768 | 841932 | 841300 | J | rpl4 | - | ko:K02926 | M00178 | Ribosomal_L4 |
| 769 | 842613 | 841972 | J | rpl3 | - | ko:K02906 | M00178,M00179 | Ribosomal_L3 |
| 782 | 852898 | 852470 | L | - | - | ko:K07492 | - | DDE_Tnp_1_2 |
| 783 | 853281 | 852895 | L | - | - | ko:K07492 | - | DUF4096 |
| 784 | 857450 | 853497 | K | rpoC2 | 2.7.7.6 | ko:K03046 | M00183 | RNA_pol_Rpb1_3,RNA_pol_Rpb1_4,RNA_pol_Rpb1_5 |
| 785 | 860866 | 857558 | K | rpoB | 2.7.7.6 | ko:K03043 | M00183 | RNA_pol_Rpb2_1,RNA_pol_Rpb2_2,RNA_pol_Rpb2_3,RNA_pol_Rpb2_45,R |
| 786 | 861737 | 860952 | L | dtd3 | - | ko:K03424 | - | TatD_DNase |
| 787 | 862166 | 861873 | J | rpsT | - | ko:K02968 | M00178 | Ribosomal_S20p |
